# Sex-specific remodeling of the tRNA epitranscriptome in Alzheimer’s disease

**DOI:** 10.64898/2026.03.31.715648

**Authors:** Marko Jörg, Lukas Walz, Marc Lander, Larissa Bessler, Sebastian Nathal, Kristine Freude, Kristina Endres, Mark Helm, Kristina Friedland

## Abstract

tRNA modifications are critical regulators of RNA stability, decoding fidelity, and cellular stress adaptation, yet their contribution to human neurodegenerative disease remains poorly understood. Beyond their established functions in translational control, emerging evidence shows that RNA modifications influence neurogenesis, neurodevelopment, neuronal function, brain-cell differentiation, and cellular plasticity. Consequently, dysregulation of these molecular processes is increasingly recognized as a mechanistic contributor to neurodegenerative disorders. Alzheimer’s disease (AD), characterized by amyloid pathology, synaptic dysfunction, and progressive neuronal loss, has recently been linked to disturbances in RNA metabolism, suggesting that alterations in the epitranscriptome may represent an underexplored dimension of AD pathophysiology. Here, we systematically profiled the tRNA epitranscriptome across cellular and animal models of AD, as well as in human postmortem brain tissue from non-demented controls and AD patients, using liquid chromatography-tandem mass spectrometry (LC-MS/MS). This method enables highly sensitive quantification of RNA modifications, with limits of detection in the low femtomole range. Across our models, we identified a conserved yet sex-specific remodeling of the tRNA modification landscape in AD. Because therapeutic options and early diagnostic tools for AD remain limited, we leveraged these findings to develop a tRNA-centered RNA-modification score that integrates both nucleobase-specific modification patterns and neuropathological disease severity into a quantitative metric. Together, our findings identify the tRNA epitranscriptome as a unifying molecular sex-specific signature of AD, linking disease pathology and sex to impaired RNA metabolism. This line of research opens a new path toward establishing early biomarkers or diagnostic tools for AD.

**Graphical abstract:** 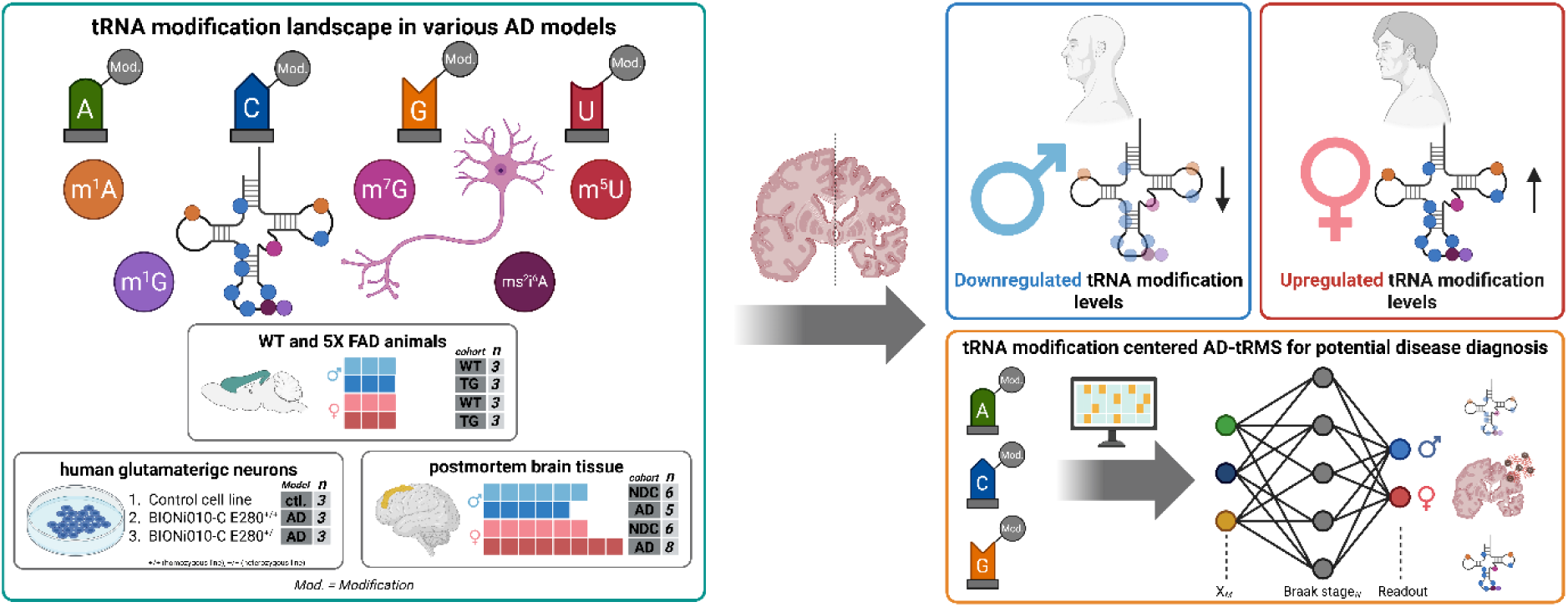

## Introduction

Alzheimer’s disease (AD) is the most common cause of dementia (60-70% of all cases), affecting more than 50 million people worldwide. It is characterized by progressive cognitive decline, synaptic loss, neuronal degeneration, and the accumulation of amyloid-beta (Aβ) plaques and tau neurofibrillary tangles (NFTs) in the brain, as well as profound mitochondrial dysfunction. Together, these pathological features promote oxidative stress, neuroinflammation, and widespread metabolic imbalance (1–8). The entorhinal cortex and hippocampus are among the earliest brain regions affected in AD, with pathology later spreading to cortical areas, likely owing to their high functional demands and susceptibility to early pathological changes (6, 9, 10). These pathological processes form a vicious cycle that progressively impairs neuronal homeostasis and causes neurodegeneration (11, 12). Despite decades of intensive research, diagnostic and therapeutic options for AD remain limited. Identifying new molecular players, prognostic biomarkers, or diagnostic tools is therefore essential to provide deeper insights into disease mechanisms, enable earlier detection, improve the effectiveness of existing therapies, and facilitate the development of novel therapeutic strategies.

Recent work has expanded the focus of AD research beyond Aβ aggregation and Tau hyperphosphorylation toward upstream regulatory mechanisms, including post-transcriptional RNA regulation and RNA modifications, thereby offering the potential to detect and understand pathogenic processes at an earlier molecular stage. RNA metabolism forms a critical interface between genomic regulation and protein synthesis, enabling cells to rapidly adapt their proteomes to physiological and pathological stimuli (13). Recent advances have highlighted the epitranscriptome, the complete set of reversible, post-transcriptional chemical modifications on RNA that regulate RNA fate, function, and gene expression, as a dynamic layer of gene regulation and a key determinant of neuronal function and survival (14). Within this framework, RNA chemical modifications have emerged as dynamic regulators of RNA fate, stability, and function, with growing implications for neuronal physiology and disease. Over 170 distinct RNA modifications have been identified across messenger RNAs (mRNAs), transfer RNAs (tRNAs), ribosomal RNAs (rRNAs), and noncoding RNAs, with the first modifications discovered as early as the 1960s (15, 16). Biochemical modifications occur across all four canonical RNA nucleosides, adenosine (A), guanosine (G), cytidine (C), and uridine (U), and collectively form a diverse and functionally essential layer of post-transcriptional regulation (13, 17–20). Among all RNA species, tRNAs represent the most extensively modified class, with cytosolic tRNAs carrying on average 13 modifications and mitochondrial tRNAs up to 5 in parallel (21, 22). These chemical modifications occur at defined structural positions within the tRNA, and their locations determine their specific functional roles, including regulation of tRNA processing, stability, subcellular localization, and overall translational efficiency (13, 22–26). In general, several distinct modified nucleosides have been described in detail, including *N*^1^-methyladenosine (m^1^A), *N*^5^-methylcytosine (m^5^C), *N*^6^-methyladenosine (m^6^A), and *N*^7^-methylguanosine (m^7^G) (27–29). Recent advances have highlighted that RNA modifications are directly implicated in neurogenesis, neurodevelopment, neuronal function, brain cell differentiation, and cellular plasticity (13, 30–38), and are increasingly associated with neurological dysfunction and neurodegenerative diseases, including AD and Parkinson’s disease (PD) (27, 39–46).

In AD, reduced levels of the mRNA modification m^6^A have been associated with enhanced Tau toxicity and altered protein expression in the 5xFAD mouse model, and age-related changes in m^6^A signatures further support a role for RNA modification in disease progression (47, 48). Beyond m^6^A, increased levels of the mRNA modification m^1^A at position 1374 of mitochondrial ND5 have been reported in AD. This TRMT10C-incorporated modification was associated with impaired mitochondrial complex I activity and respiratory dysfunction, without affecting TRMT10C-mediated tRNA modifications (m^1^A and m^1^G at position 9 of mitochondrial tRNAs) (49). These observations suggest selective remodeling of RNA modification pathways in AD and raise the possibility that additional RNA classes beyond mRNA may contribute to disease-associated epitranscriptomic alterations. Although mRNA-centered RNA modification mechanisms have been extensively investigated, the role of tRNA modifications in AD remains largely unexplored. Nevertheless, recent work demonstrates that tRNA modifications can modulate m^6^A-dependent mRNA decay pathways, providing a mechanistic link between tRNA modification status and post-transcriptional regulation (50). Given the central role of tRNAs in translation, perturbations in tRNA modification pathways represent a plausible yet largely unexplored mechanism linking AD pathology to the epitranscriptome. These pathways may also serve as early molecular sensors of neurodegenerative stress.

Here, we aimed to systematically characterize the landscape of predominantly cytosolic tRNA modifications across multiple models of AD and to determine whether these alterations represent conserved molecular features or model-specific signatures. We examined whether tRNA modification profiles exhibit consistent and sex-specific alterations across distinct AD models, whether these patterns display species-dependent differences indicative of species-specific regulatory signatures, and whether tRNA modification signatures can be integrated into a quantitative RNA modification–based metric with potential utility for diagnosis or early biomarker development. Implementing quantitative liquid chromatography–tandem mass spectrometry (LC-MS/MS), we uncovered widespread alterations in in predominantly cytosolic tRNA modifications across multiple AD models, including 5xFAD mice, high-quality human Preseniline-1 (PSEN1)^E280A induced pluripotent stem cell (hiPSC)-derived glutamatergic neurons (GlutN) carrying a PSEN1 mutation, and male and female postmortem human cortical tissue from AD patients, each analyzed in comparison with their respective matched control conditions, including non-transgenic mice, isogenic control hiPSCs-derived GlutN, and age-matched non-demented controls (NDCs). This integrative approach enabled the assessment of both conserved sex-and species-specific aspects of tRNA modification remodeling in AD.

Collectively, our findings identify sex-specific dysregulation of the tRNA epitranscriptome as a conserved molecular feature of AD. We developed a tRNA-centered RNA-modification score that integrates nucleobase-specific modification patterns with neuropathological disease severity into a quantitative metric with potential diagnostic or prognostic value from the perspective of early biomarker discovery in both male and female AD patients.

## Results

### tRNA modification profiling reveals conserved sex-specific signatures across AD models

To assess whether tRNA modification profiles are altered in AD and to determine whether such changes represent model-specific effects or conserved molecular signatures, we profiled tRNA modifications across the transgenic AD mouse model, human hiPSC-derived GlutN AD cell model, and human postmortem brain tissue. For each system, total tRNA was isolated by size-exclusion gel electrophoresis, enzymatically hydrolyzed to nucleosides, and quantified by LC-MS/MS. The LC-MS/MS method is an extremely sensitive analysis that captures the majority of known human cytoplasmic and mitochondrial tRNA modifications (Fig. 1a). In this workflow, purified tRNAs are enzymatically digested to the nucleoside level, and stable isotope-labeled internal standards (SIL-IS) are added to enable accurate quantification of individual modified nucleosides, followed by chromatographic separation and tandem mass spectrometric detection (Fig. 1b). All biological replicates were analyzed within the same LC-MS/MS batch to enable direct comparison across models, including WT mice (n = 3 males, 3 females), TG 5xFAD mice (n = 3 males, 3 females), hiPSC-derived glutamatergic neurons (control, AD PSEN1 E280 C29^+/+^, and AD PSEN1 E280 C30^+/-^; n = 3 each), and postmortem human cortical tissue (male: NDC n = 6, AD n = 5; female: NDC n = 6, AD n = 6) (Fig. 1c). We first assessed tRNA modification profiles in the 5xFAD transgenic mouse model, which develops early Aβ plaque deposition and cognitive impairments by ∼3 months of age (51, 52). LC-MS/MS profiling of cortical tissue from 12-month-old males revealed a broad reduction in nearly all quantified predominantly cytosolic tRNA modifications relative to age-matched wild-type littermates (Fig. 1d). To enable direct comparison of modification abundances within each model, we calculated Z-scores (mean = 0 for the control), with blue indicating downregulation. In contrast, female 5xFAD mice showed the opposite pattern, with most tRNA modifications upregulated relative to female wild-type controls (Fig. 1e). This pronounced divergence may indicate that alterations in tRNA modification patterns reflect sex-specific differences in AD progression. To determine whether this effect is specific to the mouse model or extends to human neuronal contexts, we next analyzed two hiPSC-derived GlutN lines harboring the PSEN1^E280A mutation. This variant represents the most common familial AD-associated PSEN1 mutation, affecting the catalytic subunit of the γ-secretase complex that regulates amyloid precursor protein (APP) processing and Aβ generation in AD (53–56). The mutation was introduced into the BIONi010-C hiPSC line derived from a healthy 18-year-old male donor using CRISPR/Cas9 editing, generating a homozygous line (BIONi010-C-29^+/+^) and a heterozygous line (BIONi010-C-30^+/-^) (53, 57). The isogenic line (UKBi004-A) was used as the control cell line (n = 3 control, n = 3 BIONi010-C-29^+/+^, n = 3 BIONi010-C-30^+/-^). LC-MS/MS analysis revealed a tRNA modification profile highly comparable to that observed in 5x FAD males, with a widespread reduction in modification levels across both PSEN1 mutant GlutN lines compared to the isogenic control hiPSC-derived GlutN (Fig. 1f). This widespread suppression suggests that sex influences multiple tRNA modification pathways rather than individual modification types, consistent with the male origin of the iPSC-derived glutamatergic neurons. Finally, to determine whether these alterations extend to human disease, we analyzed postmortem cortical tissue from male (n = 6 NDC, n = 5 AD) and female (n = 6 NDC, n = 8 AD) individuals. Consistent with our previous models, males with AD exhibited a broad reduction in tRNA modification levels relative to NDC (Fig. 1g), whereas females showed a global increase in the same modifications (Fig. 1h). To obtain an integrated overview across all systems examined, we compared the relative abundance of individual modifications across 5xFAD mice, PSEN1 mutant iPSC-derived GlutN, and human postmortem brain tissue (Fig. 1i).

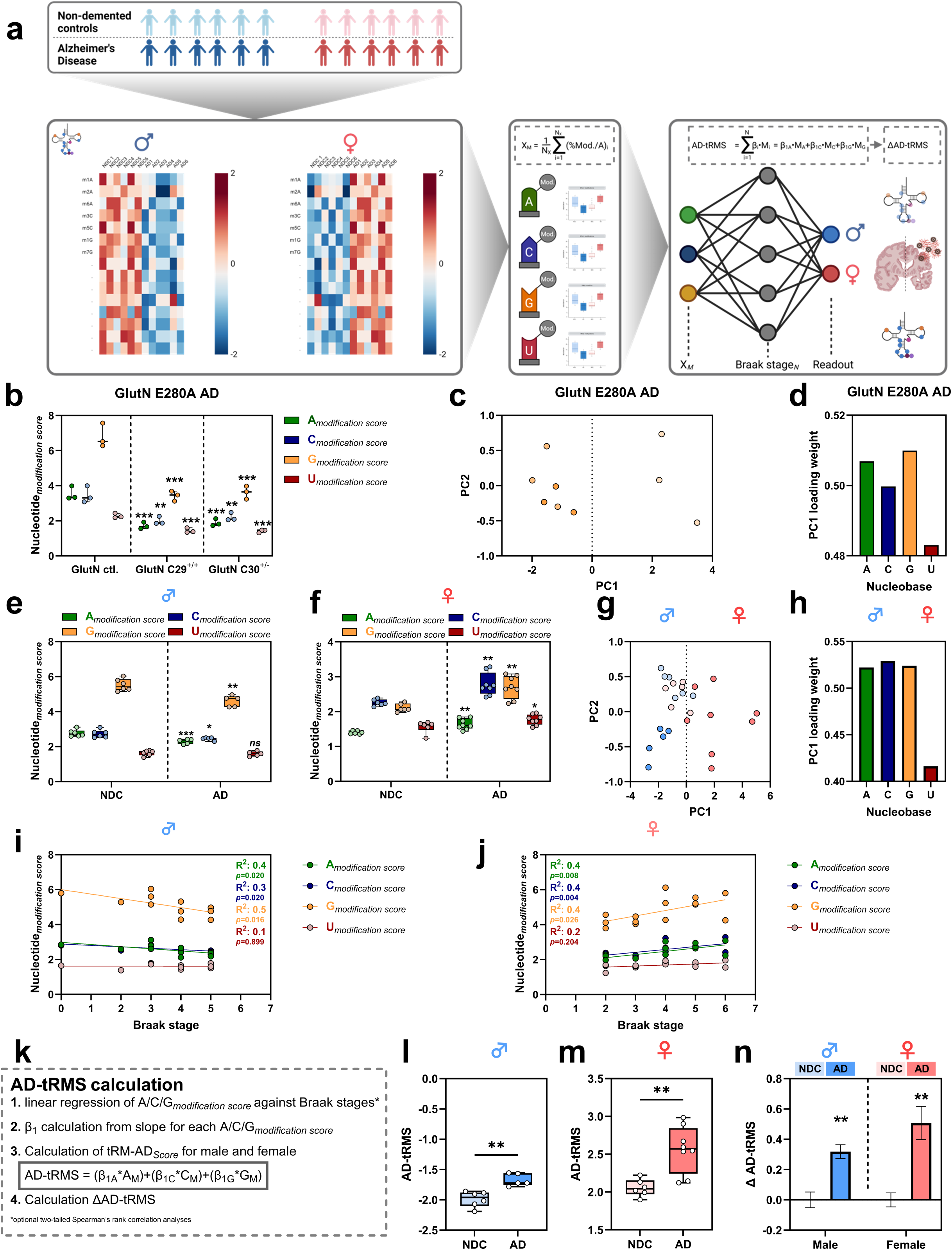

Together, these findings identify sex as a major determinant of tRNA epitranscriptomic regulation in AD, revealing a conserved, sex-specific remodeling of tRNA modifications across multiple AD models in both mouse and human systems. Such divergence may reflect distinct cellular stress responses or compensatory mechanisms that align with well-established sex differences in AD vulnerability and disease progression.

### Clustering of tRNA modification profiles distinguishes human and mouse AD models

Having established distinct sex-specific alterations in tRNA modification profiles across AD models, we next asked whether these changes are conserved across species and whether they reflect shared or model-specific molecular features of AD-associated RNA dysregulation. To address this, we performed a cross-model Pearson correlation analysis integrating LC–MS/MS–derived tRNA modification datasets from human cellular models (PSEN1^E280A iPSC-derived glutamatergic neurons), postmortem human cortical tissue, and the 5xFAD mouse model. This analysis generated a sample-to-sample similarity map, highlighting clustering of conditions based on shared modification signatures independent of absolute signal scale, with higher similarity indicated in red and lower similarity in blue revealing species-dependent segregation, with all human-derived datasets, including both cellular and brain samples, clustering tightly together. In contrast, the 5xFAD mouse samples formed a distinct, less pronounced correlated cluster (Fig. 2a). These results indicate that although tRNA modification patterns are altered across all AD systems examined, the global tRNA epitranscriptomic signatures are species-specific. Human-derived models consistently align with one another, suggesting a conserved disease-associated signature that is only partially mirrored in mouse models. To identify the specific modifications driving these separations, we performed hierarchical clustering. This unsupervised method groups samples based on similarity in individual modification abundances, thereby revealing patterns of coordinated modification changes across datasets. This analysis similarly positioned the mice samples as a distinct branch, clearly separated from all human-derived datasets (Fig. 2b). In contrast, the human models, including both in vitro and post-mortem samples, exhibited strong intercorrelation, reflecting a conserved remodeling of the tRNA modification landscape across human AD related samples. To further visualize these relationships in multidimensional space, we applied principal component analysis (PCA) and uniform manifold approximation and projection (UMAP). PCA reduces the data into orthogonal components that capture the most significant sources of variance, while UMAP preserves local and global structure to represent complex high-dimensional relationships. Both PCA and UMAP revealed a robust separation between species, with PSEN1 E280A iPSCs-derived GlutN, and human post-mortem brain samples clustering closely together, whereas mouse samples occupied a distinct region of feature space, indicating convergent molecular trajectories across human AD systems (Fig. 2c, d). To identify the tRNA modifications contributing most to the observed clustering patterns, we examined the loading weights of the principal components. The first principal component (PC1), which accounted for the most significant proportion of variance, was primarily defined by m^5^U, m^1^A, m^1^G, m^7^G, and Ψ. In contrast, the second component (PC2) was mainly driven by ms^2^i^6^A and mcm^5^U (Fig. 2e, f). We next quantified PCA-defining tRNA modifications by LC–MS/MS without dimensionality reduction or Z-score normalization to assess absolute abundances and to identify modification-specific changes across human models. Absolute quantification of selected tRNA modifications (m^1^A, m^1^G, m^7^G, ms^2^i^6^A, mcm^5^U, and Ψ) was performed using SILIS-based calibration curves, enabling precise determination of modification levels expressed as a percentage of total adenosine content (Fig. 2g, Supplementary Table S1). Using this approach, LC–MS/MS analysis revealed that the majority of the analyzed tRNA modifications were significantly reduced in PSEN1 E280A iPSC-derived glutamatergic neurons compared with their respective controls, mirroring the male-specific downregulation observed in postmortem brain tissue and consistent with the decreases identified in the previously performed normalized profiling analysis (Fig. 2h-m). In human postmortem brain tissue, we again observed pronounced sex-specific alterations: male AD patients exhibited reduced levels of the selected tRNA modifications, whereas female AD patients showed increased levels compared with NDC (Fig. 2h–m). The only exception was the mitochondria-specific tRNA modification ms²i⁶A, which was increased in PSEN1 E280A iPSC-derived cells but remained unchanged in postmortem brain tissue. This increase may reflect either a genuine upregulation of the modification or a higher mitochondrial content in glutamatergic neurons.

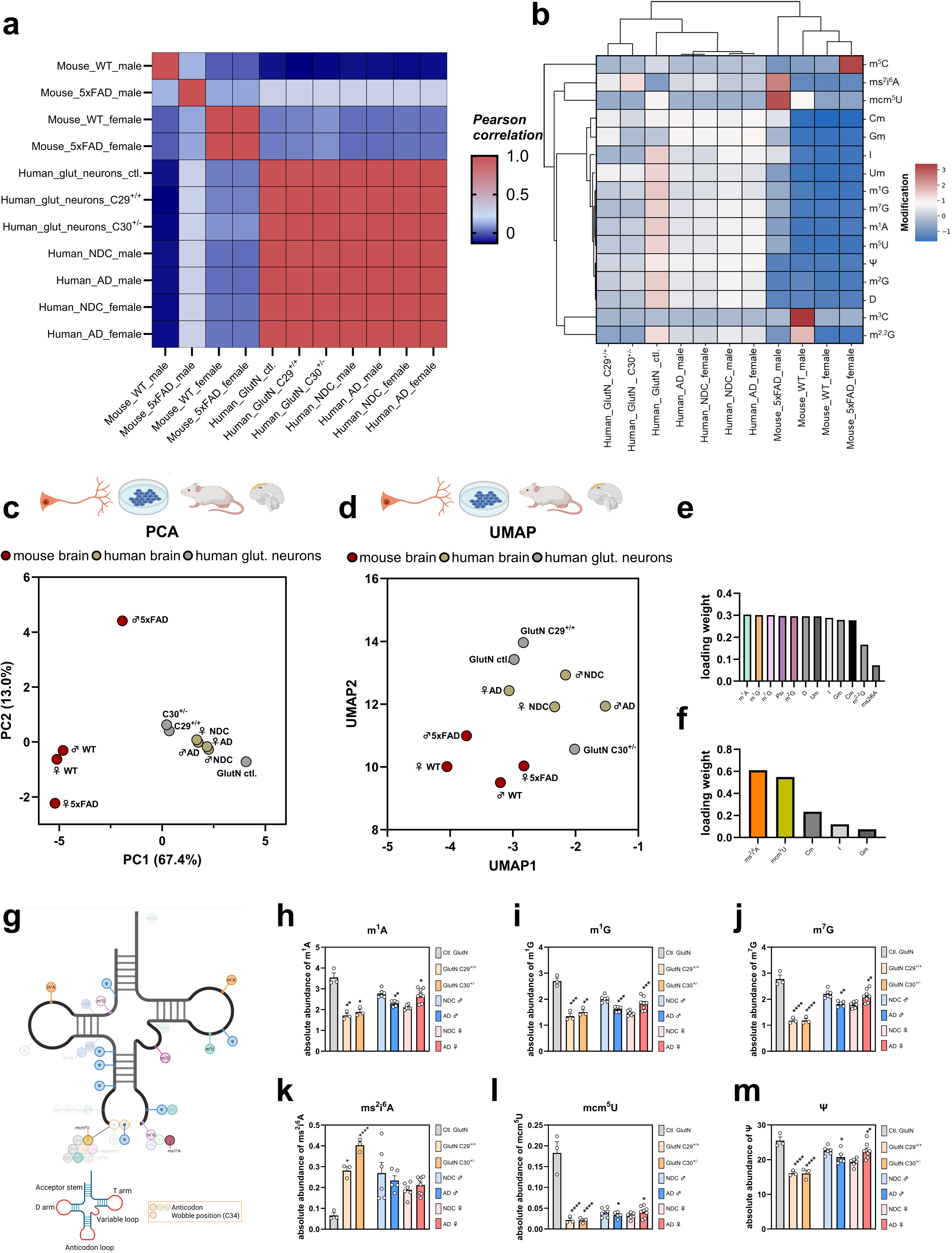

Together, these analyses reveal that tRNA modification remodeling in AD is strongly sex-dependent, with male-associated decreases and female-associated increases forming a reproducible molecular signature across human cellular and postmortem brain models. Despite overall species-specific separation between human and mouse systems, these sex-specific modification patterns were consistently preserved in human-derived datasets, indicating that sex is a dominant biological variable shaping the tRNA epitranscriptome in AD.

### Sex-specific human AD tRNA modification signatures enable development of a potential diagnostic modification score

Building on the conserved, sex-specific tRNA modification signatures observed across human postmortem brain tissue and cellular AD models, we next asked whether these coordinated epitranscriptomic changes could be integrated into a quantitative disease-associated metric for early biomarker or prognostic marker development, while accounting for the molecular and clinical complexity of AD. Our analyses are currently anchored in human brain tissue and complementary cellular systems, where disease-relevant molecular alterations are most robustly detectable. The observation that similar tRNA modification patterns emerge across distinct human models increases the likelihood that these changes reflect fundamental aspects of AD-associated RNA dysregulation and may, in principle, also manifest in other cell types and more accessible tissues, such as peripheral blood. With this in mind, and without presupposing immediate clinical applicability, we developed a nucleobase-resolved tRNA modification scoring strategy (AD-tRMS) to assess whether this composite quantitative measure tracks AD-related molecular pathology across human postmortem brain tissue and experimental models, and whether it captures biologically meaningful gradients of disease severity. In this context, the score serves as a disease-associated molecular index, providing a conceptual framework for future exploration of tRNA modification patterns in more accessible tissues and their potential utility for early detection, patient stratification, or disease monitoring in AD. We first quantified the baseline nucleobase-specific modification levels (A, C, G, and U) to establish a simplified scoring framework that could later be applied to human tissues. To ensure biological relevance and future applicability in diagnostic or biomarker development, we included only tRNA modifications with sufficient absolute abundance, defined as ≥1%. All modifications included in subsequent analyses are listed in Supplementary Table S2. For each nucleobase, we calculated an individual modification score by summing all included modifications assigned to that base and dividing this sum by the number of all integrated modifications (Fig. 3a). Applying this modification score to our human iPSC-derived GlutN model revealing a consistent reduction across all nucleobase scores in AD (Fig. 3b). Principal component analysis (PCA) demonstrated that these nucleobase-resolved scores are able to clearly separate AD model cells from isogenic control cells, indicating that coordinated shifts in tRNA modification patterns capture disease-associated variation in this model (Fig. 3c). Examination of the PC1 loading weights further revealed that A-, C-, and G-modification scores were the dominant contributors to this separation, accounting for the majority of variance along PC1. In contrast, the contribution of U-modification was markedly smaller, suggesting that uridine-linked modifications play only a minor role in driving global epitranscriptomic divergence between AD and control samples (Fig. 3d). Given the consistent performance of this score in this human cellular model, we next applied this framework to postmortem human brain tissue, a context in which substantially greater biological variability is expected. We first computed the A-, C-, G-, and U-modification scores for each postmortem sample, yielding four quantitative metrics per individual that collectively summarize nucleobase-specific modification patterns across all included modification marks. As expected, male AD patients exhibited reduced A, C, and G modification scores compared to male NDC, while U-modification showed no significant change (Fig. 3e). In contrast, female AD patients showed significantly increased A, C, and G modification relative to controls, again with only minimal effects on U modification (Fig. 3f). PCA of these methylation scores revealed clear separation not only between AD and control individuals but also between male and female AD cases (Fig. 3g). Consistent with our neuronal cell model, PC1 loading weights indicated that modifications of A, C, and G drove the clustering, whereas U modifications contributed the least (Fig. 3h). Unlike cell culture systems, human brain samples differ not only in diagnosis but also in age, sex, genetic background, and the degree of neuropathological progression. These additional sources of heterogeneity necessitate a more refined scoring strategy. To directly incorporate disease severity into the calculation, we incorporated the Braak stage, which was the only available measure of AD severity in the clinical metadata obtained from the brain bank. This established neuropathological staging system quantifies the hierarchical and region-specific spread of tau pathology in AD. By embedding the Braak stage into the modification score, we aimed to generate a more individualized, pathology-informed metric that captures biologically meaningful gradients of AD progression within human tissue (Supplementary Table S3). To formally integrate disease severity into the modification-based metric, we performed sex-stratified linear regression of each modification score against Braak stage (Fig. 3i,j). In addition, two-tailed Spearman’s rank correlation analyses revealed significant associations between Braak stage and the A-, C-, and G-modification scores in males (A*_modification score_*: *p* = 0.020; C*_modification score_*: *p* = 0.0200; G*_modification score_*: *p* = 0.016), whereas the U-modification score showed no significant correlation (*p* = 0.899). In females, Spearman’s rank correlation analysis similarly identified significant associations for the A-, C-, and G-modification scores (A*_modification score_*: *p* = 0.008; C*_modification score_*: *p* = 0.004; G*_modification score_*: *p* = 0.026), while the U-modification score again showed no significant correlation (p = 0.204). Despite the inherent heterogeneity of human postmortem tissue, the A, C, and G modification scores showed robust explanatory power in both sexes (R^2^ ≈ 0.3-0.5 in males; R^2^ ≈ 0.4 in females). In contrast, U modification exhibited only weak associations (R^2^ = 0.1-0.2), consistent with its reduced biological contribution (Fig. 3j). For each nucleobase and each sex, the regression slope (β_1_) was extracted as a weight coefficient, reflecting the contribution of that modification score to disease burden. These coefficients were incorporated into a unified composite metric, the **A**lzheimer’s **D**isease **tR**NA **M**odification **S**core (AD-tRMS): AD-tRMS = (β_1A_*A_M_)+(β_1C_*C_M_)+(β_1G_*G_M_) (Fig. 3k). The resulting AD-tRMS exhibited lower values in non-demented controls and higher values in AD cases for both men and women, as sex-specific negative and positive slopes were incorporated into the calculation (Fig. 3l,m). This approach enables the score to capture the pronounced sex-specific effects observed in tRNA modification profiles. To capture subject-specific deviation relative to controls, we additionally computed the ΔAD-tRMS, defined as the difference between each sample’s AD-tRMS and the sex-matched NDC mean. Notably, the ΔAD-tRMS revealed a similar amplitude of score effect changes in both sexes (Fig. 3n). This enables a unified readout of sex-specific AD-associated epitranscriptomic disruption. Because the AD-tRMS relies exclusively on nucleobase-specific modification measurements, it remains conceptually straightforward and analytically accessible. The computational pipeline does not require complex normalization frameworks or large reference datasets; instead, it is built directly on experimentally measured A-, C-, and G-modification levels combined with their Braak-derived weighting coefficients. Importantly, the entire workflow is technically simple. In principle, tissue extraction followed by tRNA isolation and targeted LC–MS/MS quantification would be sufficient to generate individual modification scores. From these values, the ΔAD-tRMS can be calculated with minimal computational effort. This simplicity, together with its robustness across cellular and brain tissue contexts, positions the AD-tRMS as a highly promising candidate for future diagnostic implementation, early detection of AD-related molecular changes, or patient stratification, particularly once its performance is validated in other more accessible peripheral tissues such as blood. Beyond its analytical utility, the AD-tRMS nomenclature is intentionally modular, allowing straightforward extension to tissue-specific implementations such as bAD-tRMS (blood) or cAD-tRMS (cerebrospinal fluid), thereby facilitating future translational applications.

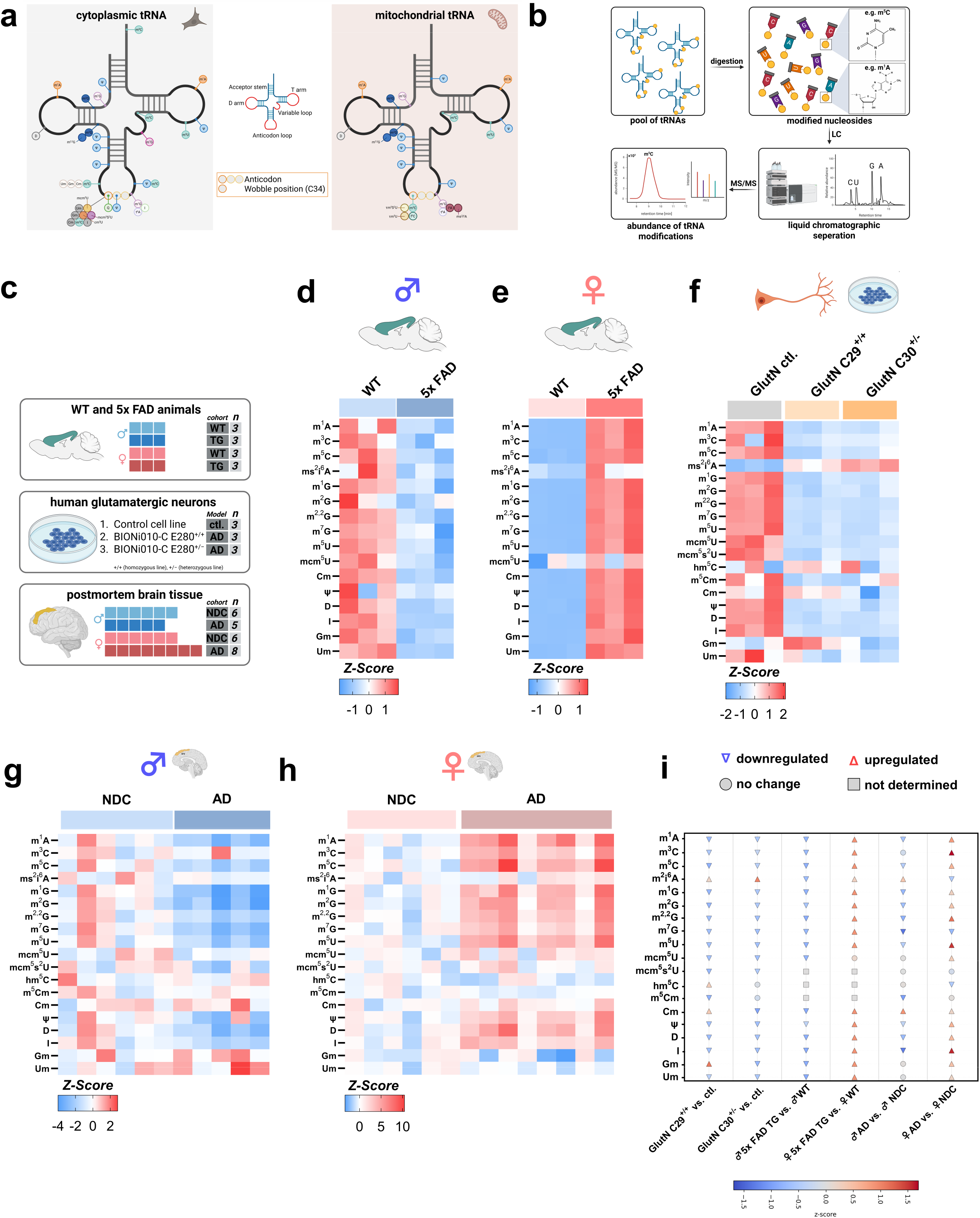

## Discussion

Our comprehensive profiling of tRNA modification landscapes across cellular, animal, and human models of AD reveals a conserved, strongly sex-specific reprogramming of the tRNA epitranscriptome. RNA modifications have long been recognized as critical regulators of RNA stability, translation, and cellular stress responses, and have gained attention both in cancer research and, more recently, as emerging contributors to neurodegenerative disorders (13, 58). Their high prevalence, evolutionary conservation, and strong context-dependent regulation position tRNA modifications as central epitranscriptomic regulators, whose disruption can profoundly affect neuronal homeostasis and vulnerability to disease (13, 58).

### Implications of sex-specific tRNA modification profiles in AD

Across both human cellular models and postmortem brain tissue, we identified a conserved subset of adenosine-, cytidine-, guanosine-, and uridine-linked tRNA modifications that exhibited consistent sex-specific dysregulation in AD. These modifications perform distinct, non-redundant functions and have been implicated in neuronal development, synaptic function, and neurological disease (13, 30–38). Consistent with our findings, prior studies have demonstrated that epitranscriptomic regulation is spatially and functionally linked to neuronal vulnerability. For example, modified RNAs, including m6A-marked transcripts, localize to synapses and regulate synaptic function and region-specific alterations in m6A mRNA levels have been reported across multiple brain regions in neurodegenerative disorders, including mild cognitive impairment and Lewy body diseases (59, 60). Moreover, analyses of small RNA methylation patterns have revealed widespread dysregulation of multiple RNA modifications in neurodegenerative contexts, aligning with the sex-dependent and multi-modification changes observed here (61). Collectively, these studies underscore that RNA modifications are not merely molecular decorations but active regulators whose disruption contributes to neurodegenerative pathology, consistent with a systematic breakdown of tRNA modification homeostasis in AD. However, most existing studies focus on the expression of writer, reader, and eraser enzymes rather than directly measuring RNA modifications and often rely on modification-specific antibodies that lack sufficient specificity for closely related nucleoside analogs (62). Such methodological limitations complicate interpretation and cross-study comparisons. By contrast, our LC–MS/MS–based approach enables direct, quantitative, and modification-resolved profiling of the tRNA epitranscriptome (63). A central finding of our study is the pronounced sex dependence of tRNA modification remodeling across models. Male-derived iPSC-derived GlutN exhibited modification losses that closely mirrored patterns observed in male AD brain tissue. Given the well-established sex-specific vulnerability in AD, where women show higher incidence and more severe progression, and the broader recognition of sex-dependent molecular differences in disease biology, our data suggest that sex-specific epitranscriptomic trajectories may contribute to these clinical disparities (64–69). Notably, Tau pathology and overall Tau burden are typically more pronounced in females, despite comparable Aβ loads, with higher p-tau181 levels and more rapid neurofibrillary tangle accumulation (70, 71). This raises the possibility that sex-dependent tRNA modification changes may intersect with Tau-associated disease mechanisms and represent an early molecular correlate of female-specific vulnerability. Contributing factors may include reduced basal autophagy in females across the lifespan, which could facilitate Tau and amyloid accumulation and secondarily influence tRNA modification profiles (71). In addition, the relative upregulation of tRNA modifications observed in females may reflect compensatory responses to estrogen loss, as estrogen supports mitochondrial function and redox homeostasis prior to menopause, with its decline in postmenopausal women (72, 73). Consistent with this, recent studies have identified regulatory links between RNA modifications and mitochondrial function, suggesting that dysregulation of RNA modification homeostasis may contribute to disease-associated mitochondrial stress (49, 74–76). Additionally, sex-specific immune and neuroinflammatory trajectories may further modulate tRNA modification dynamics, as microglial activation differs between males and females across aging, with aged female microglia exhibiting stronger inflammatory signatures than those of aged males (77, 78).

These findings underscore the necessity of sex-stratified analyses in future RNA modification studies. Within this sex-dependent framework, we also observed species-specific differences in tRNA modification dynamics. Human-derived samples clustered tightly across models, whereas WT and 5xFAD mouse samples diverged markedly. Differences in the directionality, magnitude, and sex dependence of tRNA modification remodeling highlight known limitations of mouse models in fully capturing human disease–relevant molecular regulation and may also help explain the relative resilience of mice to certain human proteinopathies (79–82).

### Validation of the tRNA modification-centered score

Given the absence of curative AD treatments, research efforts have increasingly shifted toward early diagnosis, biomarker discovery, and the identification of molecular signatures that track disease progression (83–85). Because tRNAs are extensively and diversely modified, and because these modification-dependent processes are essential for brain development, synaptic function, and neuronal health, we sought to harness nucleobase-specific tRNA modification patterns to construct a quantitative AD-associated metric. Our goal was to generate a score that reflects both the biochemical state of the tRNA epitranscriptome and the neuropathological severity of the disease. tRNAs are ideal biomarker substrates due to their multifunctional roles in translation, stress signaling, and cellular homeostasis. Analogous to established methylation-based predictors, such as Mahmoud et al.’s work on dietary methyl donors and DNA methylation patterns used as prognostic indicators in carcinogenesis and the well-known Horvath DNA methylation clock, we reasoned that tRNA modification dynamics could serve as a similarly robust readout of disease-associated molecular aging or deterioration (86, 87). The resulting AD-tRMS is designed to be analytically simple, biologically interpretable, and potentially clinically implementable. Because uridine-linked modifications contributed minimally to PC1 across all models and did not discriminate AD from control states, the U-modification score was excluded from the final composite calculation. Accordingly, the AD-tRMS was derived exclusively from A-, C-, and G-modification scores. For each sex, regression-derived coefficients (β_1_) describing the relationship between Braak stage and each nucleobase-specific modification score were used as weighting factors, yielding an individualized composite metric that integrates epitranscriptomic state with neuropathological disease severity. The AD-tRMS performed consistently across all human model systems, including hiPSC-derived GlutN and postmortem brain tissue, indicating strong cross-context robustness. While the present study establishes and validates the AD-tRMS in human brain tissue and stem cell-derived models, its applicability to peripheral tissues, including readily accessible biospecimens such as blood, remains to be determined. Given that similar tRNA modification changes are observed in both cellular models and brain tissue, it is likely that related signatures may also be detectable in peripheral blood mononuclear cells (PBMCs) or in whole blood or plasma, although this will require direct experimental validation. Nevertheless, prior studies have demonstrated that RNA modification signatures, including m^6^A on mRNA, can be robustly quantified in human blood and blood-derived immune cells, supporting the feasibility of extending RNA modification–based disease metrics to more accessible biospecimens (88, 89). However, most existing approaches focus on total RNA or mRNA and typically interrogate single modifications, most commonly m6A, thereby overlooking the complexity and functional relevance of the tRNA epitranscriptome, the most densely modified RNA class in the cell. As a consequence, coordinated, multi-site modification dynamics that may better reflect disease-associated RNA remodeling are rarely captured. By contrast, our study establishes the feasibility and biological relevance of a tRNA-centered, multi-modification framework, integrating nucleobase-specific modification patterns into a composite disease-associated metric. Nevertheless, this study is limited by tissue availability, resulting in relatively small sample sizes in the hiPSC-derived neuronal models and postmortem human brain cohorts. Future studies incorporating larger cohorts will be required to assess specificity and generalizability. If validated in larger cohorts and peripheral accessible tissues, the AD-tRMS may provide a foundation for future diagnostic, stratification, or disease-monitoring strategies in AD.

Together, our findings reveal conserved, sex-dependent tRNA modification signatures in AD, underscoring the importance of sex-specific molecular stratification for future diagnostic and individualized therapeutic approaches. Building on these insights, we introduce the AD-tRMS, which robustly captures AD-associated tRNA modification changes across human models and may facilitate future translational applications.

## Material and Methods

### Human hiPSC-derived glutamatergic neurons

Human hiPSC-derived glutamatergic neurons were provided by Kristine Freude (Faculty of Health and Medical Sciences, University of Copenhagen, Denmark) and were prepared and gene-edited as previously described by Frederiksen et al. (53). In brief, the hiPSCs lacking spontaneous differentiation were differentiated into glutamatergic neurons using a multistep directed differentiation paradigm. This workflow comprised embryoid body formation, neural rosette isolation, expansion of neural progenitor cells (NPCs), and subsequent terminal differentiation and maturation. The protocol was implemented with minor adaptations from previously established methodologies (90, 91). Neural induction medium (NIM; Supplementary Table S4) was employed during embryoid body formation and NPC specification. Following induction, NPCs were propagated on Matrigel-coated culture ware in neural expansion medium (Supplementary Table S5). For terminal differentiation, NPCs were plated at a density of 50,000 cells/cm² onto poly-L-ornithine– and laminin-coated substrates and maintained in neuronal maturation medium (NMM; Supplementary Table S6), with media replacement performed every 48 hours. For RNA isolation, 2.0 x 10^6^ cells per 10-cm dish of the C29⁺/⁺ and C30⁺/⁻ lines, and 1.5 x 10^6^ cells from isogenic control lines, were seeded for differentiation. Cells were harvested using Accutase, pelleted, and counted. Following counting, cells were washed once with PBS, detached, centrifuged, and the resulting pellet was resuspended in 1 mL TRIzol™ reagent (Thermo Fisher Scientific™, Waltham, MA, USA) and stored at -80 °C until further processing.

### Animals

Transgenic 5xFAD mice were obtained from and housed at the Translational Animal Research Center (TARC) at the University Medical Center Mainz, Germany. Animals used in this study were sex-mixed (male and female) and aged 42-55 weeks. Housing and care conditions of the animals followed the protocols described by Brandscheid et al. (92, 93). For tissue dissection, mice were anesthetized using 3% Isoflurane (Piramal Critical Care) delivered in 100% oxygen at 0.5 L/min, ensuring proper anesthetic depth by confirming loss of the interdigital reflex. Following euthanasia by a guillotine, the prefrontal cortex was carefully dissected. Animal care and experimental procedures were conducted in strict accordance with the ARRIVE guidelines and were approved by the local animal ethics committee (Landesuntersuchungsamt Rheinland-Pfalz, LUA). RNA isolation was performed as described in the section “RNA isolation from cells and tissue.”

**Table 1.** Details of LC−MS/MS detection of modified nucleosides in tRNAs.

|  | Wild-type | Transgenic animals |
| --- | --- | --- |
| Genetic background | C57BL/6J | C57BL/6J (transgen) |
| Number of samples | 6 | 6 |
| Male/Female | 3/3 | 3/3 |
| Age [weeks] |  |  |
| males | 51-63 | 51-63 |
| females | 51-63 | 51-63 |

### Postmortem cortical brain tissue

Human postmortem brain samples were obtained from the Netherlands Brain Bank (NBB, Amsterdam, The Netherlands) under project number 1341. Written informed consent for brain donation and the use of tissue and associated clinical data for research purposes was obtained from all donors or their next of kin in accordance with NBB ethical guidelines. Demented cases were clinically and neuropathologically confirmed AD patients, verified by the NBB using established biochemical and histopathological criteria. Clinical diagnoses were based on the Consortium to Establish a Registry for AD (CERAD) criteria. Non-demented control subjects exhibited no clinical or neuropathological signs of dementia at the time of death. For analysis, tissue from the superior frontal gyrus (Brodmann areas gfs 3 and gfs 4), a region relevant to AD pathology, was selected. In total, 25 brain samples were analyzed, including 14 AD cases (8 female, 6 male) and 11 non-demented controls (6 female, 5 male), with donor ages ranging from 71 to 102 years.

**Table 2:** Overview of NBB samples.

|  | Non-demented controls | Confirmed AD patients |
| --- | --- | --- |
| Number of samples | 11 | 14 |
| Male/Female | 5/6 | 6/8 |
| Age [years] (Mean ± SD) | | |
| males | 88.6 ± 10.1 | 82.5 ± 7.4 |
| females | 81.5 ± 6.3 | 84.5 ± 7.5 |
| Braak Stage (Mean ± SD) | 2.5 ± 1.0 | 4.7 ± 0.6 |

### RNA isolation from cells or tissue

TRIzol-frozen cell pellets or tissue samples were thawed at room temperature. For tissue samples, 1 mL of TRIzol^TM^ reagent (Thermo Fisher Scientific™, Waltham, MA, USA) was added, followed by homogenization using a tissue homogenizer. Chloroform (200 µL; Carl Roth, Karlsruhe, Germany) was added to each sample, which were then mixed and centrifuged at 13,000 × g for 15 min at 4 °C. The aqueous phase was transferred to a new tube, supplemented with 1 µL glycogen (Thermo Fisher Scientific™, Waltham, MA, USA) and 1 mL 100% 2-propanol (Carl Roth, Karlsruhe, Germany), and centrifuged again at 13,000 × g for 15 min at 4 °C. The supernatant was discarded, and the RNA pellet was washed with 1 mL 75% ethanol (Carl Roth, Karlsruhe, Germany), followed by centrifugation at 18,000 × g for 10 min at 4 °C. Pellets were briefly air-dried and resuspended in RNase-free water (Invitrogen, Carlsbad, CA, USA). RNA concentration and purity were determined using a NanoDrop spectrophotometer (Thermo Fisher Scientific™, Waltham, MA, USA), with purity assessed by A260/A280 and A260/A230 absorbance ratios. RNA integrity was further evaluated using an Agilent 4200 TapeStation system according to the manufacturers instructions (Agilent, Santa Clara, USA), and only samples with RNA integrity numbers (RIN^e^) ≥ 7 were included for downstream analyses (Supplementary Fig. S1).

### LC-MS/MS analysis

Total RNA samples from cell or tissue lysates were separated on a 14% TBE–Urea polyacrylamide gel (using Rotiphorese® Sequencing gel concentrate, Rotiphorese® Sequencing gel diluent and Rotiphorese® Sequencing gel buffer concentrate; Carl Roth, Karlsruhe, Germany) for 2.5h at 60 mA at 4°C. The tRNA band was visualized using SYBR™ Gold staining (1x staining solution for 30 min on a Typhoon 9400, Amersham Bioscience/GE Healthcare, Chicago, IL, USA, with 400 ms exposure time) (Thermo Fisher Scientific™, Waltham, MA, USA), excised, and re-extracted (Supplementary Fig. S2). In brief, gel slices containing the tRNA fraction were resuspended in 300 µL of 0.5 M ammonium acetate solution (Carl Roth, Karlsruhe, Germany) and frozen at −80 °C for 15 min. Samples were then shaken overnight at 15 °C at 700 rpm to facilitate RNA diffusion. The gel suspension was transferred to a spin column (Nanosep^TM^ MF centrifugal devices with Bio-Inert membrane 0.45 µm, Cytiva, Marlborough, MA, USA) and centrifuged at 2,000 × g for 5 min. The filtrate was supplemented with 1 µL glycogen (Thermo Fisher Scientific^TM^, Waltham, MA, USA) and 700 µL of prechilled (−80 °C) 100% ethanol (Carl Roth, Karlsruhe, Germany), then incubated at −80 °C for 45 min. RNA was pelleted by centrifugation at 18,000 × g for 90 min at 4 °C, and the supernatant was discarded. The resulting pellet was air-dried and resuspended in nuclease-free water (Thermo Fisher Scientific, Waltham, MA, USA). RNA concentration and purity were determined spectrophotometrically using a NanoDrop instrument (Thermo Fisher Scientific, Waltham, MA, USA).

For LC-MS/MS analysis purified tRNA (1000 ng) was digested to the nucleoside level according to Thüring et al. using 0.6 U nuclease P1 (from *Penicillium citrinum*), 0.2 U snake venom phosphodiesterase (from *Crotalus adamanteus*, Worthington), 2 U FastAP (Thermo Scientific), 10 U Benzonase (Sigma-Aldrich), 200 ng pentostatin (Sigma-Aldrich), and 25 mM ammonium acetate buffer (pH 7.5; Sigma-Aldrich) (63). Digestions were performed overnight at 37 °C. Each sample was analyzed in at least three biological replicates. For each LC-MS/MS injection, 500 ng tRNA samples were mixed with a ^13^C-labeled internal standard (ISTD) to allow absolute quantification and calibration. Measurements were performed on an Agilent 1260 Infinity II LC system (Agilent, Santa Clara, USA) coupled to an Agilent 6460 Triple Quadrupole mass spectrometer. Chromatographic separation was achieved on a Synergi™ Fusion RP C18 column (S-4 µm, 80 Å, 250 × 2.0 mm I.D.; Phenomenex, Torrance, USA) maintained at 35 °C. The LC gradient (NUCS5 method) used phase A (acetonitrile, Honeywell Riedel-de Haën) from 0% to 8% over 10 min, followed by phase B (5 mM NH_4_OAc, pH 5.3), then a 10 min equilibration at a flow rate of 0.35 mL min⁻¹. UV detection was performed using a multiple-wavelength detector (MWD) at 254 nm with an attenuation of 1000 mAU and a sampling rate of 2.5 Hz. Mass spectrometric detection was conducted via an ESI source operated in positive ion mode under the following conditions: capillary current = 5400 nA, gas temperature = 350 °C, sheath gas temperature = 345 °C, sheath gas flow = 10.0 L min⁻¹, gas flow = 8.0 L min⁻^1^, nebulizer pressure = 50 psi, corona voltage = 0 V. Multiple reaction monitoring (MRM) windows of ±3 min were applied around optimized retention times for the following modified nucleosides and their ^13^C-labeled standards: m^1^A, m^3^C, m^5^C, ms^2^i^6^A, m^1^G, m^2^G, m^2,2^G, m^7^G, m^5^U, mcm^5^U, mcm^5^s^2^U, hm^5^C, Cm, Ψ, D, I, Gm, and Um (abbreviations according to MODOMICS). The acquisition parameters are given in Table 1. Examples of LC–ESI–MRM–MS/MS raw chromatograms are shown in Supplementary Fig. S3.

### LC-MS/MS data analysis

Chromatograms and mass spectra were analyzed using Agilent MassHunter Qualitative Analysis software. UV chromatograms recorded at 254 nm were evaluated first, and peak areas corresponding to the canonical nucleosides (C, U, G, and A) were manually integrated. UV peak areas derived from stable isotope-labeled internal standards (SIL-IS) were subtracted from the total sample peak areas to obtain corrected values. Calibration was performed using SIL-IS calibration solutions containing equal amounts of labeled nucleosides, prepared as described by Thüring et al. (63). For quantitative analysis, both unlabeled (^12^C) and labeled (^13^C) peak areas were extracted, and the absolute amount of each modified ucleoside was calculated using a response factor-based equation. All RNA samples were spiked with identical amounts of SIL-IS, and the calculated quantities of modified nucleosides were normalized to the corresponding canonical nucleosides.

### Bioinformatics and data analysis

Hierarchical clustering analysis was performed to identify patterns in RNA modification profiles across experimental groups. The dataset comprised 16 RNA modification markers measured across 13 experimental conditions, including mouse models (wild-type and 5xFAD, both male and female), human hiPSC-derived GlutN (isogenic control, C29^+/+^, and C30^+/-^), and human postmortem samples (NDC and AD, both male and female). Data preprocessing included Z-score standardization (mean = 0, standard deviation = 1) to normalize for differences in measurement scales across RNA modification markers. Hierarchical clustering was performed using the average linkage method with Euclidean distance metric for both rows (markers) and columns (experimental groups). PCA was conducted on the transposed and standardized dataset to identify major sources of variance across experimental groups. The data matrix was transposed such that experimental groups served as observations and RNA modification markers as features. Prior to PCA, data were standardized using Z-score normalization to ensure equal weighting of all markers regardless of their original measurement scales. PCA was performed using singular value decomposition (SVD) with mean centering. The first three principal components were retained based on the Kaiser criterion and cumulative variance explained. Component loadings were calculated to identify the contribution of each RNA modification marker to the principal components, with the top contributors to PC1 being m^5^U, m^1^G, m^1^A, m^7^G, and Um modifications. UMAP analysis was performed as a complementary nonlinear dimensionality-reduction technique to capture local neighborhood structures not revealed by linear PCA. The same standardized and transposed dataset used for PCA was subjected to UMAP analysis using the following parameters: n_neighbors=5 (to focus on local structure), min_dist=0.1 (to allow for tight clustering), n_components=2 (for 2D visualization), and random_state=42 (for reproducibility). The UMAP algorithm constructs a high-dimensional graph representation of the data based on local neighborhoods, then optimizes a low-dimensional embedding that preserves the topological structure. This approach is particularly effective at revealing nonlinear relationships and local clustering patterns that may be obscured by linear dimensionality reduction methods. All analyses were performed in Python 3.11 using the following packages: pandas (v1.5.3) for data manipulation and analysis, NumPy (v1.24.4) for numerical computations and array operations, scikit-learn (v1.6.1) for PCA, standardization, and preprocessing utilities, SciPy (v1.11.3) for hierarchical clustering algorithms and linkage methods, UMAP-learn (v0.5.9) for non-linear dimensioality reduction and manifold learning, openpyxl (v3.1.2) for Excel file generation and multi-sheet workbook creation. Hierarchical clustering dendrograms and heatmaps were generated using seaborn’s clustermap function, with custom formatting parameters optimized for publication-quality output. PCA visualizations included both individual component plots and cumulative variance plots to assess the number of components required to explain the data structure. UMAP visualizations were created with consistent color schemes and marker sizes to facilitate direct comparison with PCA results.

### Visualization of results and statistics

All graphs were generated using Prism GraphPad 10.6.1 (GraphPad Software Inc., La Colla, CA, USA) or Python 3.11. Statistical analyses were performed in Prism GraphPad 10.6.1 (GraphPad Software Inc., La Colla, CA, USA)using Student’s unpaired t-test for comparisons between two groups or one-way ANOVA followed by Tukey’s post hoc test for multiple-group comparisons, depending on the experimental design and group size. Correlations were assessed using two-tailed Spearman’s rank correlation. Data are presented as mean + standard error of the mean (SEM) unless otherwise stated. A significance threshold of p < 0.05 was applied for all statistical tests.

## Supporting information

Main figure legends

## Illustrations

All illustrations were created using BioRender, including publication license (https://BioRender.com).

## Data availability

The data underlying this article are available in the article and in its online supplementary material.

## Author contributions

M.J. and K.F. conceptualized and designed the project. M.J. performed the majority of the experimental work. K.E. provided the animals. M.J. and L.W. carried out dissections. M.J., S.N., and L.W carried out RNA isolation. M.J., M.J., M.L. and L.B. performed LC-MS/MS measurements. K.Fr. provided human hiPSC-derived glutamatergic neurons and performed culturing, differentiation and sample collection. L.W., M.J., carried out bioinformatic data analysis, M.J., M.H., and K.F discussed results altogether. M.J. and K.F wrote the manuscript. All authors reviewed and edited the manuscript.

## Supplementary data

Supplementary Data are available at Nature Communications online.

## Funding

Deutsche Forschungsgemeinschaft (DFG, German Research Foundation) [TRR-319 TP A05 to M.H., Project Id 439 669 440].

## Conflict of interest

Mark Helm is a consultant for Moderna Inc.

---

**Table 1:** Overview of mouse samples.

| Dynamic MRM |  |  |  |  |  |
| --- | --- | --- | --- | --- | --- |
| Compound name | Precursor ion | Product ion | Fragmentor | Retention time [min] | Polarity |
| m<sup>1</sup>A | 282.0 | 150.0 | 80.0 | 5.3 | Positive |
| m<sup>1</sup>A 13C | 293.0 | 156.0 | 80.0 | 5.3 | Positive |
| m<sup>3</sup>C | 258.0 | 126.0 | 80.0 | 4.6 | Positive |
| m<sup>3</sup>C 13C | 268.0 | 131.0 | 80.0 | 4.6 | Positive |
| m<sup>5</sup>C | 258.0 | 126.0 | 75.0 | 8.6 | Positive |
| m<sup>5</sup>C 13C | 268.0 | 131.0 | 75.0 | 8.6 | Positive |
| ms<sup>2</sup>i<sup>6</sup>A | 382.0 | 250.0 | 70.0 | 38.0 | Positive |
| ms<sup>2</sup>i<sup>6</sup>A 13C | 398.0 | 261.0 | 70.0 | 38.0 | Positive |
| m<sup>1</sup>G | 298.0 | 166.0 | 85.0 | 13.6 | Positive |
| m<sup>1</sup>G 13C | 309.0 | 172.0 | 85.0 | 13.6 | Positive |
| m<sup>2</sup>G | 298.0 | 166.0 | 85.0 | 13.6 | Positive |
| m<sup>2</sup>G 13C | 309.0 | 172.0 | 85.0 | 13.6 | Positive |
| m<sup>2,2</sup>G | 312.0 | 180.0 | 95.0 | 17.3 | Positive |
| m<sup>2,2</sup>G 13C | 324.0 | 187.0 | 95.0 | 17.3 | Positive |
| m<sup>7</sup>G | 298.0 | 166.0 | 80.0 | 7.4 | Positive |
| m<sup>7</sup>G 13C | 309.0 | 172.0 | 80.0 | 7.4 | Positive |
| m<sup>5</sup>U | 259.0 | 127.0 | 70.0 | 11.0 | Positive |
| m<sup>5</sup>U 13C | 269.0 | 132.0 | 70.0 | 11.0 | Positive |
| mcm<sup>5</sup>U | 317.0 | 185.0 | 90.0 | 14.1 | Positive |
| mcm<sup>5</sup>U 13C | 329.0 | 192.0 | 90.0 | 14.1 | Positive |
| mcm<sup>5</sup>s<sup>2</sup>U | 333.0 | 201.0 | 80.0 | 19.7 | Positive |
| mcm<sup>5</sup>s<sup>2</sup>U 13C | 345.0 | 208.0 | 80.0 | 19.7 | Positive |
| hm<sup>5</sup>C | 274.0 | 142.0 | 90.0 | 7.3 | Positive |
| hm<sup>5</sup>C 13C | 284.0 | 147.0 | 90.0 | 7.4 | Positive |
| m<sup>5</sup>Cm | 272.0 | 126.0 | 80.0 | 12.8 | Positive |
| m<sup>5</sup>Cm 13C | 283.0 | 131.0 | 80.0 | 12.8 | Positive |
| Cm | 258.0 | 112.0 | 80.0 | 9.5 | Positive |
| Cm 13C | 268.0 | 116.0 | 80.0 | 9.5 | Positive |
| Ψ | 245.0 | 209.0 | 85.0 | 4.2 | Positive |
| Ψ 13C | 254.0 | 218.0 | 85.0 | 4.2 | Positive |
| D | 247.0 | 115.0 | 90.0 | 4.0 | Positive |
| D 13C | 256.0 | 119.0 | 90.0 | 4.0 | Positive |
| I | 269.0 | 137.0 | 75.0 | 10.1 | Positive |
| I 13C | 279.0 | 142.0 | 75.0 | 10.1 | Positive |
| Gm | 298.0 | 152.0 | 85.0 | 13.9 | Positive |
| Gm 13C | 309.0 | 157.0 | 85.0 | 13.9 | Positive |
| Um | 259.0 | 113.0 | 75.0 | 12.2 | Positive |
| Um 13C | 269.0 | 117.0 | 75.0 | 12.2 | Positive |

