## Supplement for "Sex-specific remodeling of the tRNA epitranscriptome in Alzheimer’s disease"


### Supplementary Informations

**Abbreviations**

| Abbreviation | Name |
|---|---|
| A | Adenosine |
| AD | Alzheimer's Disease |
| AD-tRMS | Alzheimer's Disease tRNA Modification Score |
| APP | Amyloid precursor protein |
| Aβ | Amyloid β |
| C | Cytosine |
| G | Guanosine |
| Glutamatergic neurons | GlutN |
| hiPSC | high-quality induced pluripotent stem cell |
| LC-MS/MS | liquid chromatography-tandem mass spectrometry |
| mRNA | messenger RNA |
| NDC | Non-demented control |
| NFT | neurofibrillary tangle |
| PC1 | Principal component 1 |
| PC2 | Principal component 2 |
| PCA | Principal component analysis |
| PSEN1 | Preseniline-1 |
| rRNA | ribosomal RNA |
| SIL-IS | stable isotope-labeled internal standards |
| U | Uridine |
| UMAP | uniform manifold approximation and projection |

**RNA Modifications:**

| Abbreviation | Name* |
|---|---|
| $m^1A$ | $N^1$-methyladenosine |
| $m^3C$ | 3-methylcytidine |
| $m^5C$ | 5-methylcytosine |
| $ms^2i^6A$ | 2-methylthio-N6-isopentenyladenosine |
| $m^1G$ | 1-methylguanosine |
| $m^2G$ | $N^2$-methylguanosine |
| $m^{2,2}G$ | $N^2$,$N^2$-dimethylguanosine |
| $m^7G$ | 7-methylguanosine |
| $m^5U$ | 5-methyluridine |
| $mcm^5U$ | 5-methoxycarbonylmethyluridine |
| $mcm^5s^2U$ | 5-methoxycarbonylmethyl-2-thiouridine |
| $hm^5C$ | 5-hydroxymethylcytidine |
| $m^5Cm$ | 5,2'-O-dimethylcytidine |
| Cm | 2'-O-methylcytidine |
| ψ | Pseudouridine |
| D | Dihydrouridine |
| I | Inosine |
| Gm | 2'-O-methylguanosine |
| Um | 2'-O-methyluridine |

*names according to MODOMICS

### Supplementary Figures

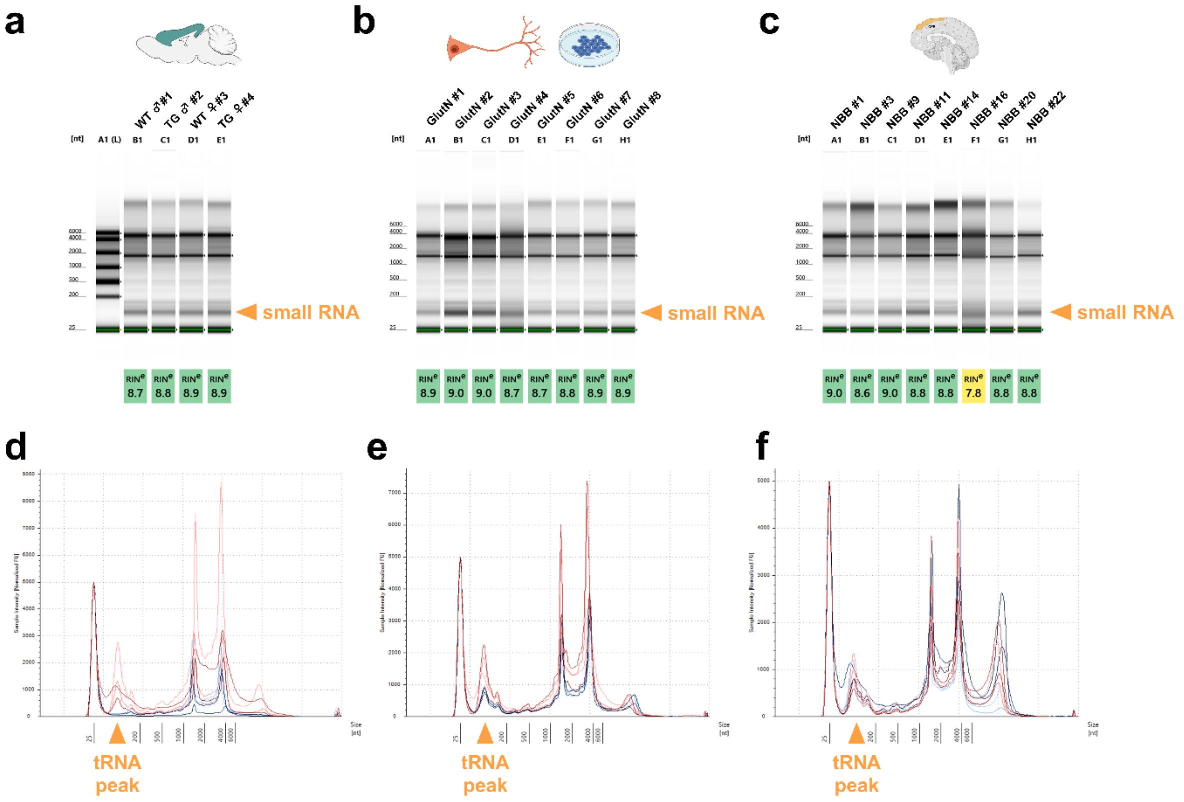


**Supplementary Fig. S1. RNA sample quality control.** Total RNA from all samples was initially assessed by absorbance ratios (A260/A280 and A260/A230) as described in Materials and Methods. Representative samples from mouse tissue, hiPSC-derived glutamatergic neurons (GlutN), and postmortem human cortical brain tissue cohorts were additionally analyzed by automated capillary electrophoresis using an Agilent 4200 TapeStation system. (a-c) Automated gel electrophoresis profiles for mouse samples (a), hiPSC-derived glutamatergic neurons (b), and postmortem cortical brain tissue (c). All analyzed samples exhibited RNA integrity equivalent (RINe) values > 7, indicating high RNA quality without detectable degradation. (d–f) Representative TapeStation electropherograms of total RNA from mouse (d), GlutN (e), and postmortem brain samples (f). The tRNA peak (~75–100 nt) is highlighted in orange. Prominent peaks at approximately 75–100 nt, ~1,900 nt, and ~5,000 nt correspond to tRNA, 18S rRNA, and 28S rRNA, respectively, confirming intact RNA integrity. In (d), only red traces correspond to samples included in this study.

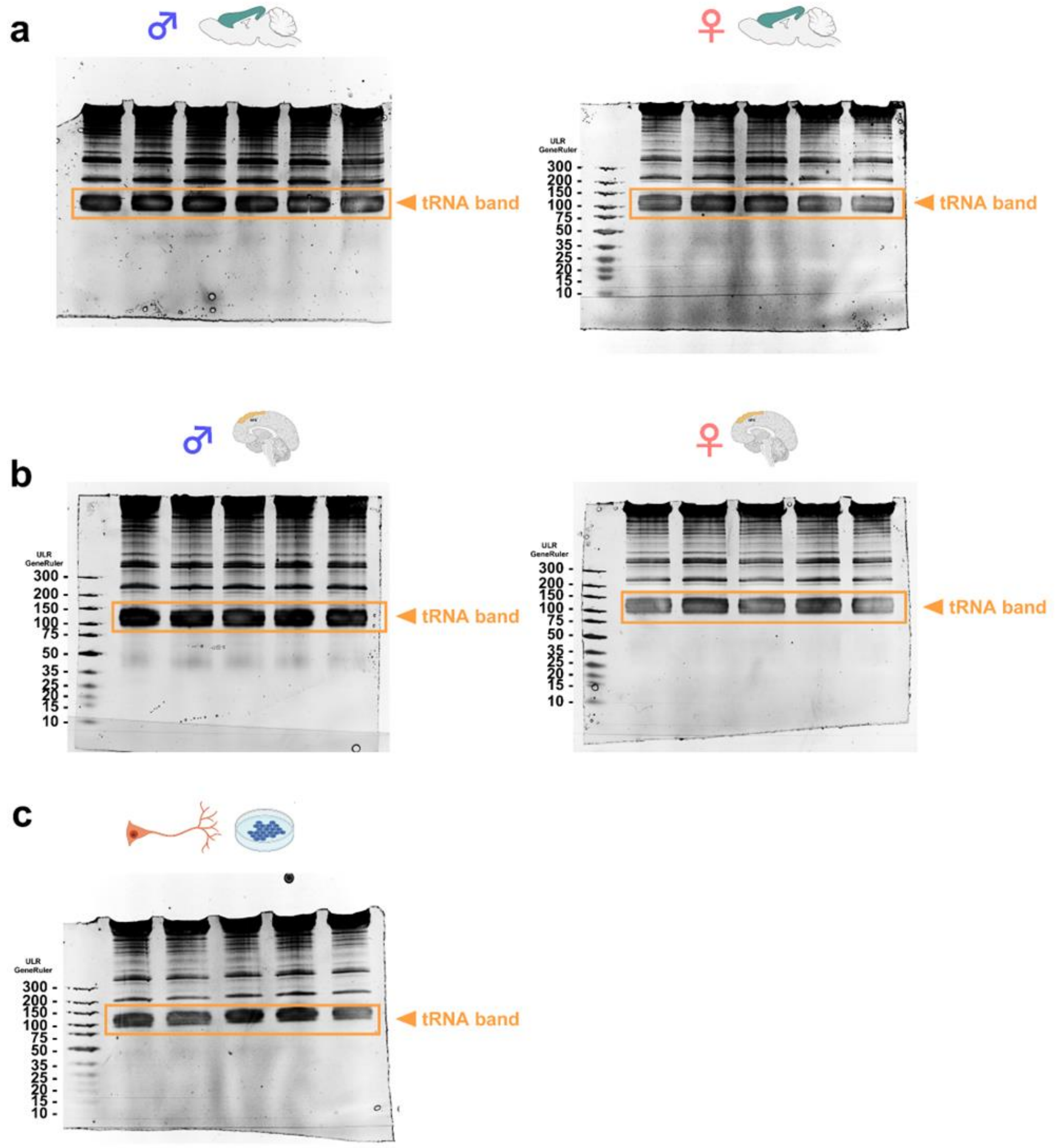


**Supplementary Fig. S2. Representative PAGE gels for tRNA isolation.** Quality-controlled total RNA was separated by denaturing polyacrylamide gel electrophoresis (PAGE) as described in the Materials and Methods section. (a-c) Representative gels showing male and female wild-type and transgenic mouse samples (a), male and female non-demented controls (NDC) and Alzheimer's disease (AD) postmortem cortical brain samples (b), and hiPSC-derived glutamatergic neurons (GlutN) (c). For each lane, 30 µg of total RNA was loaded. The orange box indicates the tRNA band, which was excised for subsequent tRNA purification and LC-MS/MS analysis.

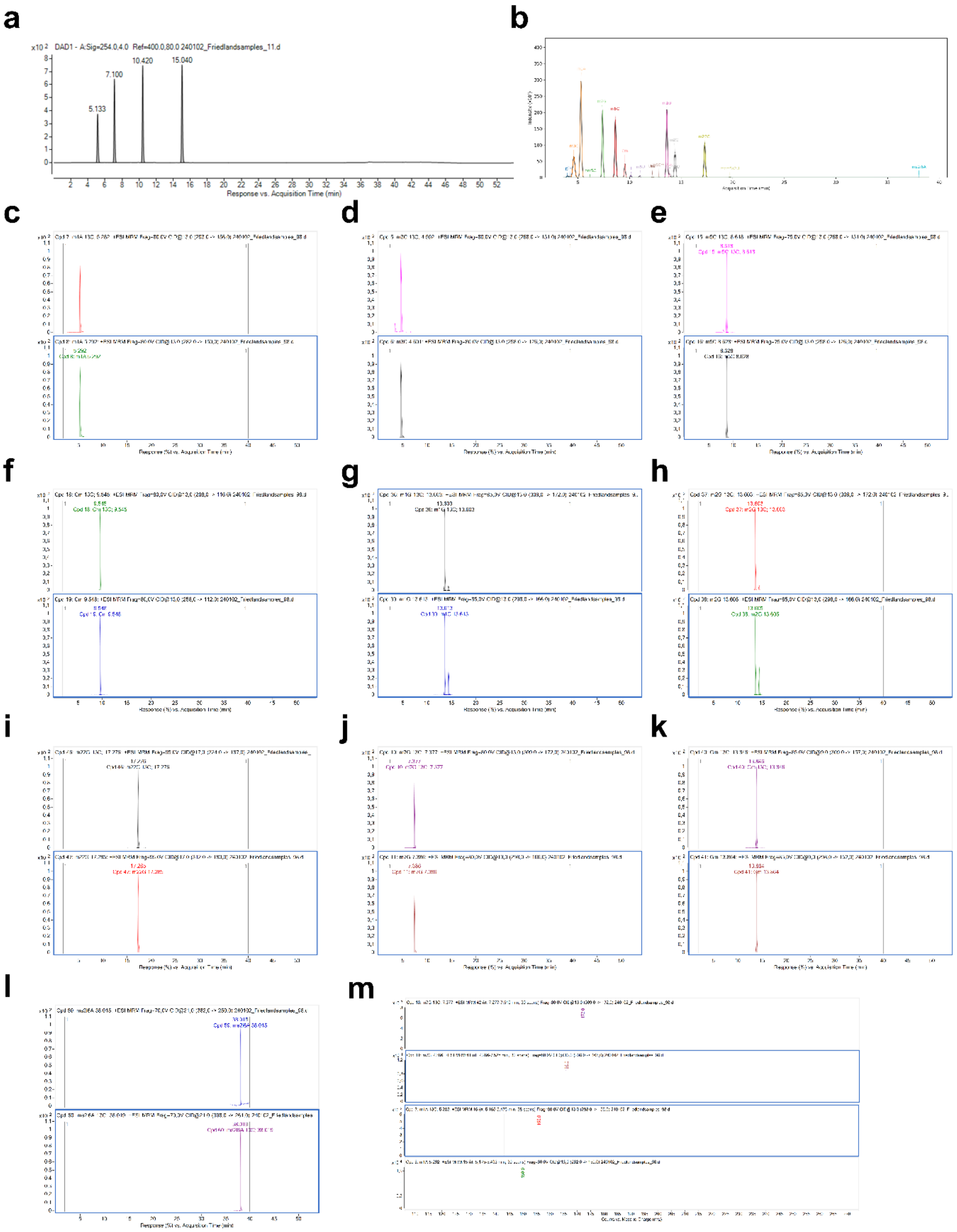


**Supplementary Fig. S3. LC–ESI–MRM–MS/MS validation of CUGA and modified ribonucleosides**. (a) Representative chromatographic profile of CUGA showing a distinct and well-resolved peak at the expected retention time, confirming successful separation under the applied LC conditions. (b) Baseline-resolved chromatographic separation of modified ribonucleosides used for quantitative analysis. (c–l) Representative multiple reaction monitoring (MRM) chromatograms for individual modified ribonucleosides. For each analyte, one trace shows the 13C standard, and the other trace displays the corresponding endogenous compound, demonstrating co-elution and analytical specificity. (m) Representative MS/MS fragmentation spectra of selected RNA modifications confirming compound identity based on characteristic precursor-to-product ion transitions. Chromatographic separation was performed by liquid chromatography coupled to electrospray ionization tandem mass spectrometry (LC–ESI–MS/MS) in MRM mode with optimized fragmentor voltages and collision energies for each analyte.

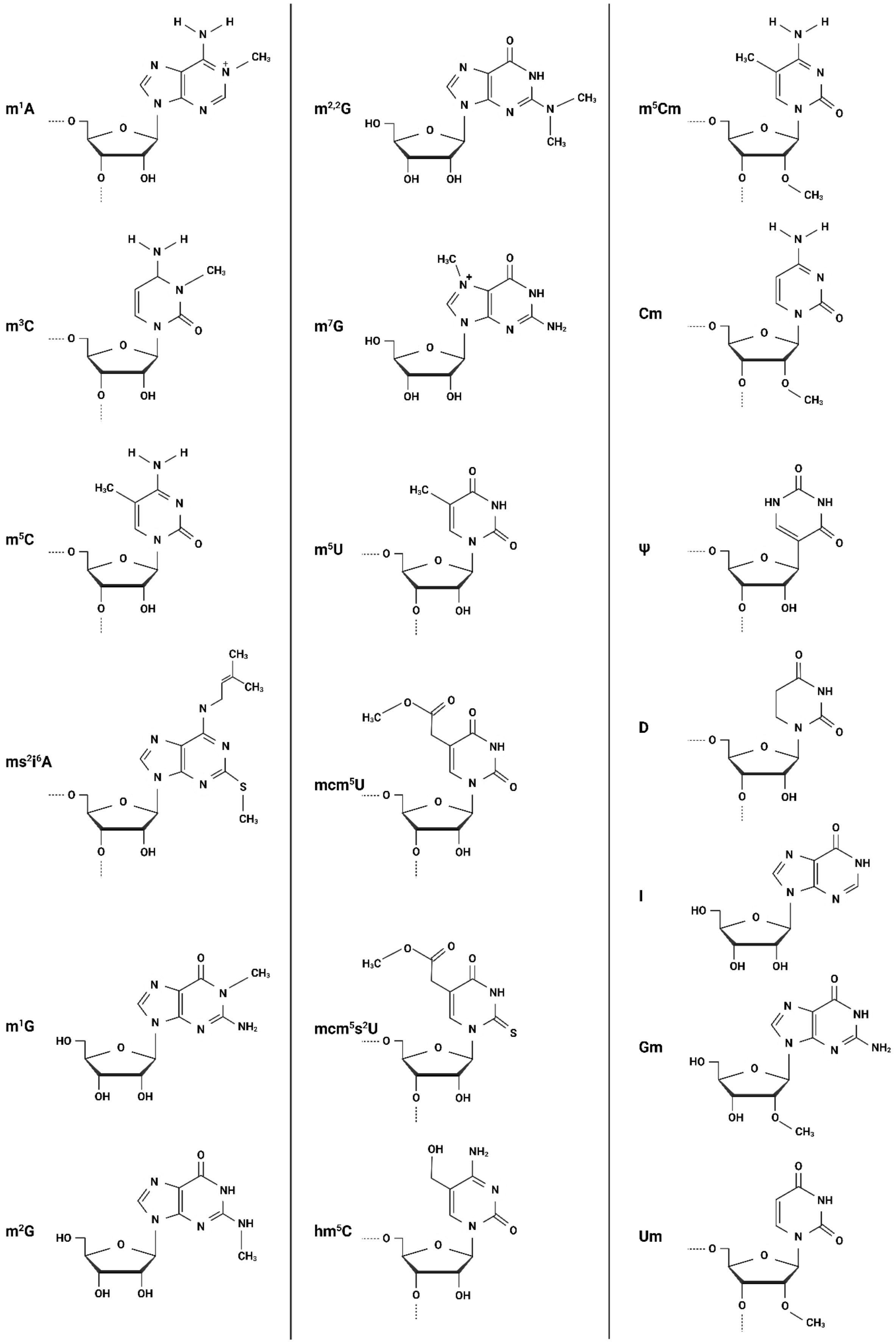


**Supplementary Figure S4.** Structures of RNA modifications measured in Alzheimer's disease models.

### Supplementary tables

Supplementary Table S1. Absolute quantification of RNA modifications in tRNA

| Model | background | $m^1A$ | $m^3C$ | $m^5C$ | $ms^2i^6A$ | $m^1G$ | $m^2G$ | $m^{2,2}G$ | $m^7G$ | $m^5U$ | $mcm^5U$ | $mcm^5s^2U$ | $hm^5C$ | $m^5Cm$ | Cm | ψ | D | I | Gm | Um |
|---|---|---|---|---|---|---|---|---|---|---|---|---|---|---|---|---|---|---|---|---|
| GlutN #1 | control | 3.300 | 1.576 | 7.357 | 0.052 | 2.523 | 21.902 | 4.391 | 2.653 | 2.480 | 0.195 | 0.430 | 0.001 | 0.007 | 0.970 | 24.611 | 23.511 | 1.477 | 1.105 | 3.077 |
| GlutN #2 | control | 3.338 | 0.340 | 7.988 | 0.091 | 2.662 | 21.069 | 4.130 | 2.584 | 2.460 | 0.223 | 0.413 | 0.002 | 0.004 | 0.912 | 24.110 | 23.549 | 1.26 | 0.924 | 3.831 |
| GlutN #3 | control | 3.990 | 1.975 | 8.913 | 0.055 | 2.914 | 25.467 | 5.160 | 3.087 | 2.965 | 0.131 | 0.336 | 0.001 | 0.008 | 1.106 | 27.704 | 27.628 | 1.573 | 1.225 | 2.582 |
| GlutN #4 | $C29^{+/+}$ | 1.666 | 0.462 | 4.106 | 0.298 | 1.25 | 11.844 | 1.400 | 1.147 | 1.381 | 0.016 | 0.214 | 0.002 | 0.002 | 0.995 | 15.530 | 10.409 | 0.620 | 1.712 | 2.847 |
| GlutN #5 | $C29^{+/+}$ | 1.554 | 0.513 | 4.161 | 0.244 | 1.231 | 10.430 | 1.235 | 1.095 | 1.260 | 0.018 | 0.205 | 0.002 | 0.003 | 0.942 | 15.814 | 9.388 | 0.535 | 1.627 | 2.269 |
| GlutN #6 | $C29^{+/+}$ | 1.922 | 0.674 | 5.131 | 0.302 | 1.560 | 12.899 | 1.424 | 1.294 | 1.505 | 0.031 | 0.264 | 0.001 | 0.002 | 0.972 | 16.928 | 11.860 | 0.554 | 1.280 | 2.024 |
| GlutN #7 | $C30^{+/-}$ | 1.818 | 0.634 | 4.732 | 0.409 | 1.432 | 13.224 | 1.293 | 1.130 | 1.295 | 0.013 | 0.240 | 0.002 | 0.002 | 0.903 | 17.144 | 10.441 | 0.525 | 1.182 | 2.563 |
| GlutN #8 | $C30^{+/-}$ | 1.731 | 0.633 | 4.838 | 0.370 | 1.409 | 11.65 | 1.257 | 1.097 | 1.303 | 0.023 | 0.235 | 0.001 | 0.004 | 0.725 | 14.228 | 10.512 | 0.547 | 0.765 | 2.127 |
| GlutN #9 | $C30^{+/-}$ | 2.122 | 0.753 | 5.809 | 0.431 | 1.674 | 14.484 | 1.476 | 1.332 | 1.556 | 0.027 | 0.263 | 0.001 | 0.006 | 0.872 | 16.772 | 12.556 | 0.609 | 0.902 | 2.230 |
| NBB #1 | ♀ AD | 2.642 | 1.132 | 6.036 | 0.173 | 1.851 | 16.674 | 3.355 | 2.111 | 2.174 | 0.058 | 0.298 | 0.00 | 0.005 | 1.096 | 21.815 | 17.367 | 1.281 | 1.791 | 2.134 |
| NBB #2 | ♀ AD | 2.721 | 1.140 | 6.155 | 0.170 | 1.683 | 18.746 | 3.510 | 2.195 | 2.225 | 0.026 | 0.420 | 0.00 | 0.004 | 1.010 | 22.482 | 19.441 | 1.177 | 1.609 | 2.419 |
| NBB #3 | ♀ AD | 3.081 | 1.266 | 7.435 | 0.248 | 2.124 | 18.897 | 3.837 | 2.401 | 2.518 | 0.048 | 0.507 | 0.001 | 0.005 | 1.151 | 24.350 | 22.369 | 1.205 | 1.769 | 2.360 |
| NBB #4 | ♀ AD | 2.234 | 0.946 | 5.249 | 0.15 | 1.54 | 13.254 | 2.824 | 1.824 | 1.901 | 0.022 | 0.214 | 0.001 | 0.004 | 1.061 | 19.826 | 15.245 | 0.889 | 1.813 | 2.019 |
| NBB #5 | ♀ AD | 2.649 | 1.042 | 5.929 | 0.253 | 1.858 | 17.434 | 3.281 | 2.023 | 2.121 | 0.055 | 0.281 | 0.001 | 0.003 | 1.054 | 22.649 | 18.231 | 1.031 | 1.572 | 2.450 |
| NBB #6 | ♀ AD | 2.948 | 1.08 | 5.926 | 0.238 | 1.879 | 17.177 | 3.365 | 2.111 | 2.181 | 0.038 | 0.291 | 0.00 | 0.005 | 1.028 | 22.478 | 19.001 | 1.051 | 1.531 | 1.739 |
| NBB #7 | ♀ AD | 2.283 | 0.881 | 5.500 | 0.214 | 1.552 | 13.277 | 2.826 | 1.861 | 1.884 | 0.044 | 0.233 | 0.001 | 0.006 | 1.016 | 20.613 | 15.353 | 0.980 | 1.845 | 2.485 |
| NBB #8 | ♀ AD | 3.043 | 1.287 | 7.228 | 0.249 | 2.209 | 20.194 | 4.058 | 2.501 | 2.614 | 0.041 | 0.357 | 0.00 | 0.006 | 1.180 | 26.805 | 20.709 | 1.258 | 1.633 | 2.148 |
| NBB #9 | ♂ AD | 2.410 | 0.954 | 5.35 | 0.282 | 1.709 | 15.459 | 3.015 | 1.906 | 1.916 | 0.023 | 0.340 | 0.001 | 0.004 | 1.066 | 21.533 | 17.597 | 1.004 | 1.705 | 2.306 |
| NBB #10 | ♂ AD | 2.420 | 0.971 | 5.541 | 0.29 | 1.712 | 16.615 | 3.085 | 1.937 | 1.908 | 0.039 | 0.292 | 0.001 | 0.004 | 1.027 | 19.742 | 17.683 | 0.921 | 1.498 | 1.847 |
| NBB #11 | ♂ AD | 2.120 | 1.538 | 4.898 | 0.174 | 1.497 | 13.905 | 2.607 | 1.697 | 1.678 | 0.032 | 0.273 | 0.00 | 0.006 | 1.015 | 18.975 | 15.081 | 0.828 | 1.707 | 1.916 |
| NBB #12 | ♂ AD | 2.362 | 0.902 | 5.458 | 0.213 | 1.681 | 15.588 | 2.926 | 1.841 | 1.862 | 0.034 | 0.340 | 0.001 | 0.005 | 1.152 | 23.931 | 16.634 | 0.883 | 1.815 | 2.716 |
| NBB #13 | ♂ AD | 2.184 | 0.923 | 5.007 | 0.211 | 1.514 | 13.873 | 2.76 | 1.727 | 1.688 | 0.041 | 0.292 | 0.001 | 0.004 | 0.973 | 19.553 | 15.347 | 0.836 | 1.490 | 2.413 |
| NBB #14 | ♀ NDC | 2.279 | 0.984 | 5.094 | 0.200 | 1.602 | 14.715 | 2.898 | 1.796 | 1.736 | 0.029 | 0.311 | 0.001 | 0.005 | 1.089 | 20.630 | 15.809 | 0.879 | 1.797 | 2.451 |
| NBB #15 | ♀ NDC | 2.064 | 0.875 | 4.864 | 0.219 | 1.36 | 12.786 | 2.671 | 1.666 | 1.720 | 0.038 | 0.224 | 0.001 | 0.007 | 0.951 | 19.705 | 13.973 | 0.770 | 1.698 | 2.261 |
| NBB #16 | ♀ NDC | 2.216 | 0.784 | 5.214 | 0.13 | 1.462 | 12.941 | 2.676 | 1.824 | 1.817 | 0.035 | 0.307 | 0.001 | 0.010 | 1.007 | 20.042 | 14.231 | 0.908 | 1.683 | 2.532 |

| | | | | | | | | | | | | | | | | | | | | |
|---|---|---|---|---|---|---|---|---|---|---|---|---|---|---|---|---|---|---|---|---|
| NBB #17 | ♀ NDC | 1.975 | 0.844 | 4.721 | 0.229 | 1.380 | 12.013 | 2.269 | 1.629 | 1.536 | 0.030 | 0.253 | 0.001 | 0.001 | 1.038 | 17.311 | 13.295 | 0.707 | 1.684 | 1.562 |
| NBB #18 | ♀ NDC | 2.066 | 0.908 | 4.810 | 0.177 | 1.557 | 14.245 | 2.798 | 1.906 | 1.589 | 0.019 | 0.357 | 0.001 | 0.005 | 1.042 | 19.411 | 15.383 | 0.978 | 1.803 | 2.631 |
| NBB #19 | ♀ NDC | 2.066 | 0.874 | 4.540 | 0.182 | 1.443 | 13.041 | 2.589 | 1.930 | 1.501 | 0.040 | 0.175 | 0.00 | 0.008 | 1.004 | 18.717 | 15.223 | 0.891 | 1.807 | 2.899 |
| NBB #20 | ♂ NDC | 2.662 | 1.060 | 5.718 | 0.360 | 1.858 | 17.929 | 3.337 | 2.046 | 2.064 | 0.027 | 0.353 | 0.002 | 0.006 | 0.882 | 20.939 | 18.261 | 1.142 | 1.407 | 1.756 |
| NBB #21 | ♂ NDC | 3.118 | 1.302 | 7.010 | 0.194 | 2.226 | 20.066 | 3.941 | 2.476 | 2.33 | 0.037 | 0.317 | 0.001 | 0.008 | 1.007 | 24.498 | 22.186 | 1.295 | 1.47 | 2.085 |
| NBB #22 | ♂ NDC | 2.848 | 1.225 | 6.137 | 0.151 | 2.039 | 19.154 | 3.703 | 2.268 | 2.168 | 0.027 | 0.296 | 0.001 | 0.006 | 1.049 | 22.431 | 20.779 | 1.163 | 1.82 | 1.912 |
| NBB #23 | ♂ NDC | 2.596 | 0.622 | 5.873 | 0.460 | 2.013 | 18.011 | 2.912 | 2.128 | 1.85 | 0.053 | 0.342 | 0.00 | 0.001 | 1.05 | 21.811 | 18.973 | 0.896 | 1.363 | 1.589 |
| NBB #24 | ♂ NDC | 2.792 | 0.880 | 6.531 | 0.302 | 2.053 | 18.886 | 3.333 | 2.240 | 2.109 | 0.042 | 0.341 | 0.001 | 0.002 | 1.067 | 23.086 | 19.652 | 1.067 | 1.501 | 2.271 |
| NBB #25 | ♂ NDC | 2.610 | 1.154 | 5.735 | 0.148 | 1.903 | 16.748 | 3.518 | 2.088 | 2.109 | 0.051 | 0.348 | 0.001 | 0.005 | 0.962 | 22.757 | 18.831 | 1.033 | 1.505 | 2.197 |

Table S2. tRNA modifications contributing to modification scores

| $X_{\text{modification score}}$ | contributing tRNA modification (abs. quant. levels > 1%) |
|---|---|
| $A_{\text{modification score}}$ | $m^1A$ |
| $C_{\text{modification score}}$ | $m^3C$, $m^5C$, Gm |
| $G_{\text{modification score}}$ | $m^1G$, $m^2G$, $m^{2,2}G$, $m^7G$, Gm |
| $U_{\text{modification score}}$ | $m^5U$, Um |

Table S3. tRNA modification scores

| **Diagnosis** | **Brain region** | **Sex** | **$A_{\text{modification score}}$** | **$C_{\text{modification score}}$** | **$G_{\text{modification score}}$** |
|---|---|---|---|---|---|
| AD | prefrontal cortex | F | 2.641819408 | 2.754840145 | 5.15661135 |
| AD | prefrontal cortex | F | 2.72062635 | 2.768225165 | 5.548521877 |
| AD | prefrontal cortex | F | 3.080583112 | 3.283969548 | 5.805671252 |
| AD | prefrontal cortex | F | 2.233982226 | 2.418557575 | 4.250687901 |
| AD | prefrontal cortex | F | 2.648703520 | 2.675098047 | 5.233720126 |
| AD | prefrontal cortex | F | 2.948423098 | 2.677823392 | 5.212355438 |
| AD | prefrontal cortex | F | 2.282627759 | 2.465282867 | 4.272162054 |
| AD | prefrontal cortex | F | 3.043152262 | 3.231719009 | 6.118819669 |
| AD | prefrontal cortex | M | 2.409854100 | 2.456809921 | 4.758862785 |
| AD | prefrontal cortex | M | 2.420263306 | 2.513067664 | 4.969584110 |
| AD | prefrontal cortex | M | 2.119793743 | 2.483618203 | 4.282463514 |
| AD | prefrontal cortex | M | 2.362063614 | 2.503742201 | 4.770231828 |
| AD | prefrontal cortex | M | 2.184419055 | 2.301101399 | 4.272828092 |
| Control | prefrontal cortex | F | 2.278926295 | 2.388972817 | 4.561657634 |
| Control | prefrontal cortex | F | 2.064085041 | 2.230132023 | 4.036088828 |
| Control | prefrontal cortex | F | 2.215955600 | 2.335099377 | 4.117082820 |
| Control | prefrontal cortex | F | 1.975306389 | 2.200938018 | 3.794765394 |
| Control | prefrontal cortex | F | 2.065852056 | 2.253458360 | 4.461690939 |
| Control | prefrontal cortex | F | 2.065861604 | 2.139463900 | 4.162086280 |
| Control | prefrontal cortex | M | 2.662003733 | 2.553106199 | 5.315416716 |
| Control | prefrontal cortex | M | 3.117894134 | 3.106206509 | 6.035528716 |
| Control | prefrontal cortex | M | 2.847853764 | 2.803524686 | 5.796578537 |
| Control | prefrontal cortex | M | 2.596489015 | 2.515224215 | 5.285395272 |
| Control | prefrontal cortex | M | 2.791889129 | 2.826083375 | 5.602732637 |
| Control | prefrontal cortex | M | 2.610218618 | 2.617221880 | 5.152330126 |

Supplementary Table S4. NIM (Neural Induction Medium) for glutamatergic neurons differentiation

| **Compound** | **Company and Catalog number** | **Final concentration** |
|---|---|---|
| Neurobasal medium | Gibco, 21103049 | 47% |
| DMEM-F12 | Gibco, 11330032 | 47% |
| Penicillin-Streptomycin | Sigma Aldrich, P0781 | 1% |
| GlutaMax™ (100X) | Gibco, 35050-038 | 1% |
| Non-essential Amino Acid (100×) | Sigma Aldrich, M7145 | 1% |
| N-2 Supplement (100X) | Gibco, 17502001 | 1% |
| B-27™ Supplement (50X) | Gibco, 15440584 | 2% |
| SB431542 | StemMACS, 130-106-543 | 10 µM |
| LDN193189 | Sigma Aldrich, SML0559 | 0.1µM |

Supplementary Table S5. NEM (Neural Expansion Medium) for glutamatergic neurons differentiation

| Compound | Company and Catalog number | Final concentration |
|---|---|---|
| Neurobasal medium | Gibco, 21103049 | 47% |
| DMEM-F12 | Gibco, 11330032 | 47% |
| Penicillin-Streptomycin | Sigma Aldrich, P0781 | 1% |
| GlutaMax™ (100X) | Gibco, 35050-038 | 1% |
| Non-essential Amino Acid (100×) | Sigma Aldrich, M7145 | 1% |
| N-2 Supplement (100X) | Gibco, 17502001 | 1% |
| B-27™ Supplement (50X) | Gibco, 15440584 | 2% |
| bFGF2 | Peprotech, 100-18B | 20 ng/ml |
| bEGF | Prospec, CYT-217 | 20 ng/ml |

Supplementary Table S6. NMM (Neural Maturation Medium) for glutamatergic neurons differentiation

| Compound | Company and Catalog number | Final concentration |
|---|---|---|
| Neurobasal medium | Gibco, 21103049 | 47% |
| DMEM-F12 | Gibco, 11330032 | 47% |
| Penicillin-Streptomycin | Sigma Aldrich, P0781 | 1% |
| GlutaMax™ (100X) | Gibco, 35050-038 | 1% |
| Non-essential Amino Acid (100×) | Sigma Aldrich, M7145 | 1% |
| N-2 Supplement (100X) | Gibco, 17502001 | 1% |
| B-27™ Supplement (50X) | Gibco, 15440584 | 2% |
| L-ascrobic acid | HelloBio, HB1238 | 200 μM |
| db-cAMP | Sigma Aldrich, D0260 | 50 μM |
| GDNF | Peprotech, 100-18B | 10 ng/ml |
| BDNF | Prospec, CYT-217 | 20 ng/ml |
