## Supplementary material for "Sex-specific remodeling of the tRNA epitranscriptome in Alzheimer’s disease": Main figure legends

**Figure 1. tRNA modification profiling identifies sex-specific AD-associated signatures.** a) Schematic representation of cytoplasmic and mitochondrial tRNA modifications. b) Workflow for global tRNA modification profiling. Total tRNA was isolated by size-based gel purification, enzymatically digested to nucleosides, and quantified by LC–MS/MS. c) Overview of AD models analyzed, including PSEN1$^{E280A}$ hiPSC-derived glutamatergic neurons, 5xFAD mice (male and female), and postmortem human cortical tissue from NDC and AD patients (male and female). d) Male 5xFAD mice display widespread loss of tRNA modifications relative to male wild-type littermates. e) Female 5xFAD mice show the opposite effect, with global upregulation of tRNA modifications compared with female wild-type controls. f) PSEN1$^{E280A}$ hiPSC-derived glutamatergic neurons (heterozygous and homozygous) exhibit a global reduction in tRNA modifications compared with the isogenic control line. g–h) Postmortem cortical tissue displays the same sex-dependent divergence: males with AD show reduced tRNA modification levels, whereas females exhibit increased levels relative to age-matched NDC. i) Integrated comparison across all models reveals a conserved sex-specific AD-associated remodeling of the tRNA modification landscape, with consistent downregulation in human male-derived AD samples, and upregulation in female mouse and human AD samples. *n* = 3-8 per group.

**Figure 2. Clustering of tRNA modification profiles separates human and mouse AD models.** a) Cross-model Pearson correlation matrix of LC–MS/MS-derived tRNA modification datasets from $PSEN1^{E280A}$ hiPSC-derived glutamatergic neurons, postmortem human cortex, and 5xFAD mouse cortex. Human-derived samples cluster together, whereas 5xFAD samples form a distinct cluster, indicating species-specific modification signatures. b) Hierarchical clustering of tRNA modification profiles further separates 5xFAD samples from all human-derived datasets, reflecting conserved remodeling of the tRNA modification landscape across human AD models. c–d) PCA and UMAP visualization of tRNA modification profiles show strong convergence among human models and distinct separation from 5xFAD samples. e–f) PCA loading plots identify the modified nucleosides $m^1A$, $m^1G$, $m^7G$, $m^5U$, Ψ, $ms^2i^6A$, and $mcm^5U$ as primary contributors to variance across models. g) Representative tRNA modifications for absolute quantification. h-m) Absolute quantification using SIL-IS calibration curves confirms significant decreases in tRNA modification abundances across human models, except $ms^2i^6A$, which is increased in $PSEN1^{E280A}$ hiPSC-derived glutamatergic neurons. $n$ = 3-8 per group. Statistical analysis was performed using ordinary one-way ANOVA followed by Dunnett's multiple comparison test or students' unpaired t-test (**$p<0.01$, ***$p<0.001$, ****$p<0.0001$).

**Figure 3. Development and validation of a nucleobase-resolved tRNA modification score for AD diagnosis (AD-tRMS).** a) Schematic of the nucleobase-resolved modification scoring strategy. For each nucleobase (A, C, G, U), individual tRNA modifications with ≥1% absolute abundance were summed and normalized to the number of contributing modifications with incorporating braak stages for weighting disease burden. b) Application of the modification score in human hiPSC-derived glutamatergic neurons. AD hiPSC-derived GlutNs show reduced nucleobase modification scores (b), clear separation by PCA (c), and dominant contributions of A-, C-, and G-modification to PC1 (d). e-f) Nucleobase-resolved modification scores in postmortem human brain tissue reveal sex-specific effects: reduced A-, C-, and G-modification in male AD cases (e) and increased A-, C-, and G-modification in female AD cases (f), with minimal changes in U modification. g–h) PCA of postmortem samples separates AD from controls and distinguishes male and female AD cases, driven primarily by A-, C-, and G-modification scores (h). i–j) Sex-stratified linear regression of nucleobase modification scores against Braak stage demonstrates strong associations for A-, C-, and G-modification, but weak explanatory power for U-modification. Correlation was analyzed using Spearman's rank correlation. k) Construction of the composite Alzheimer's Disease tRNA Modification Score (AD-tRMS) by weighting A-, C-, and G-modification scores with Braak-derived regression coefficients ($\beta_1$). l-m) The AD-tRMS discriminates AD cases from non-demented controls in both males and females. q) ΔAD-tRMS, calculated relative to sex-matched control means. *n* = 3-8 per group. Statistical analysis was performed using ordinary one-way ANOVA followed by Dunnett's multiple comparison test or students unpaired t-test, (***p*<0.01, ****p*<0.001, *****p*<0.0001).
